# Killer whale genomes reveal a complex history of recurrent admixture and vicariance

**DOI:** 10.1101/520718

**Authors:** Andrew D. Foote, Michael D. Martin, Marie Louis, George Pacheco, Kelly M. Robertson, Mikkel-Holger S. Sinding, Ana R. Amaral, Robin W. Baird, C. Scott Baker, Lisa Ballance, Jay Barlow, Andrew Brownlow, Tim Collins, Rochelle Constantine, Willy Dabin, Luciano Dalla Rosa, Nicholas J. Davison, John W. Durban, Ruth Esteban, Steven H. Ferguson, Tim Gerrodette, Christophe Guinet, M. Bradley Hanson, Wayne Hoggard, Cory J. D. Matthews, Filipa I. P. Samarra, Renaud de Stephanis, Sara B. Tavares, Paul Tixier, John A. Totterdell, Paul Wade, M. Thomas P. Gilbert, Jochen B.W. Wolf, Phillip A. Morin

## Abstract

Reconstruction of the demographic and evolutionary history of populations assuming a consensus tree-like relationship can mask more complex scenarios, which are prevalent in nature. An emerging genomic toolset, which has been most comprehensively harnessed in the reconstruction of human evolutionary history, enables molecular ecologists to elucidate complex population histories. Killer whales have limited extrinsic barriers to dispersal and have radiated globally, and are therefore a good candidate model for the application of such tools. Here, we analyse a global dataset of killer whale genomes in a rare attempt to elucidate global population structure in a non-human species. We identify a pattern of genetic homogenisation at lower latitudes and the greatest differentiation at high latitudes, even between currently sympatric lineages. The processes underlying the major axis of structure include high drift at the edge of species’ range, likely associated with founder effects and allelic surfing during post-glacial range expansion. Divergence between Antarctic and non-Antarctic lineages is further driven by ancestry segments with up to four-fold older coalescence time than the genome-wide average; relicts of a previous vicariance during an earlier glacial cycle. Our study further underpins that episodic gene flow is ubiquitous in natural populations, and can occur across great distances and after substantial periods of isolation between populations. Thus, understanding the evolutionary history of a species requires comprehensive geographic sampling and genome-wide data to sample the variation in ancestry within individuals.

## 1 INTRODUCTION

Genetic divergence of isolated populations can be interrupted by episodic gene flow during periods of spatial contact, which can erode genetic differences between populations (**Durand et al., 2009**; **Gompert et al., 2010**), rescue small isolated populations (**Frankham, 2015**), and maintain standing genetic variation that can act as a substrate for future adaptation (**Brawand et al., 2014**; **Meier et al., 2017**). The geographic context of ancestral spatial contact is difficult to elucidate from the contemporary distribution of modern populations (**Pickrell & Reich, 2014**; **Foote, 2018; Peñalba et al., 2018**), especially in marine species, which can have dynamic ranges due to the low energetic cost of movement and few physical barriers in the oceans (**Gagnaire et al., 2015**; **Kelley et al., 2016**). Additionally, ancestral episodes of admixture may have occurred via now-extinct ‘archaic’ populations (**Racimo et al., 2015**) or sister species (**Fraïsse et al., 2015**), further complicating the inference of the biogeographic history of ancestry components from the spatial distribution of contemporary populations. However, periods of admixture leave genomic signatures that can be used to infer the direction, extent and timing of ancestral gene flow (**Patterson et al., 2012**; **Sousa & Hey, 2013**; **Racimo et al., 2015**; **Duranton et al., 2018**). Divergence-with-gene-flow is often studied at local scales, but could influence global genetic structure and variation through connected networks of populations (**Novembre et al., 2008; Booth Jones et al., 2018**).

Killer whales have a global distribution rivalling that of modern humans, yet they can exhibit fine-scale geographic variation in ecology and morphology (**Ford et al., 1998**; **Durban et al., 2017**), reflecting variation in their demographic and evolutionary history (**Hoelzel et al., 2007**; **Morin et al., 2015**; **Foote et al., 2016**). The best-studied ecotypes are the mammal-eating ‘*transients*’ and fish-eating ‘*residents*’ found in partial sympatry throughout the coastal waters of the North Pacific (**Ford et al., 1998**; **Saulitis et al., 2000**; **Matkin et al., 2007**; **Filatova et al., 2015**). Four decades of field observations have found that *residents* and *transients* are socially isolated and genetically differentiated across their geographic range (**Hoelzel & Dover, 1991**; **Barrett-Lennard, 2000**; **Ford, Ellis & Balcomb, 2000**; **Hoelzel et al., 2007**; **Morin et al., 2010**; **Parsons et al., 2013**; **Filatova et al., 2015**). There has been a contentious debate regarding whether the formation of these two ecotypes was initiated in sympatry (**Moura et al., 2015**), or results from secondary contact of two distinct lineages (**Foote et al., 2011**; **Foote & Morin, 2015, 2016**).

In the waters around the Antarctic continent, killer whales have diversified into distinct morphs, partially overlapping in their ranges (**Pitman & Ensor, 2003**; **Pitman et al., 2007**; **Durban et al., 2017**). These include *type B1*, which is commonly observed hunting seals in the pack-ice; *type B2*, which has been observed foraging in open water for penguins; and *type C*, which is most commonly observed in the dense pack-ice, and is thought to primarily feed on fish (**Pitman & Ensor, 2003**; **Pitman & Durban, 2010, 2012**; **Durban et al., 2017**). Perhaps surprisingly given their highly distinct morphological forms (**Pitman & Ensor, 2003**; **Pitman et al., 2007**; **Durban et al., 2017**), the Antarctic types are inferred from previous genomic analyses to have diversified from a recent shared ancestral lineage following an extended genetic bottleneck (**Morin et al., 2015**; **Foote et al., 2016**). However, the reconstruction of the evolutionary relationships among ecotypes, and how these relate to a more globally distributed dataset, has proved challenging due to incomplete lineage sorting and admixture, and a paucity of genomic data from a wider geographic distribution (**Foote & Morin, 2016**).

Here we fill this gap by providing 27 additional genomes to a global dataset totalling 47 genomes and trace population history of separation and admixture. First, we describe the global patterns of biodiversity, then we focus on the history of well characterised ecotypes of the North Pacific and Antarctica to test between the opposing hypotheses of simple history of vicariance versus a more complex (and previously hidden) history involving ancient secondary contact between divergent lineages.

## 2 MATERIALS AND METHODS

### 2.1 Dataset

Genome sequences were generated from 27 individuals that best represented the known global geographic, genetic and morphological diversity of this species (**Figure 1a**). For a subset of analyses, we further included 20 previously sequenced genomes (**Supporting Information Table S1**): four additional genomes each from the North Pacific *transient* and *resident* ecotypes, and Antarctica types *B1*, *B2* and *C* (**Foote et al., 2016**), and 10 RAD-seq generated genotypes (**Moura et al., 2015**). In addition, we sequenced an outgroup sample of a long-finned pilot whale (*Globicephala melas*) from a mass stranding at Ratmanoff beach, Kerguelen island in the Southern Ocean and included sequencing reads of the bottlenose dolphin (*Tursiops truncatus*, Short Read Archive accession code SRX200685).

**Figure 1.**
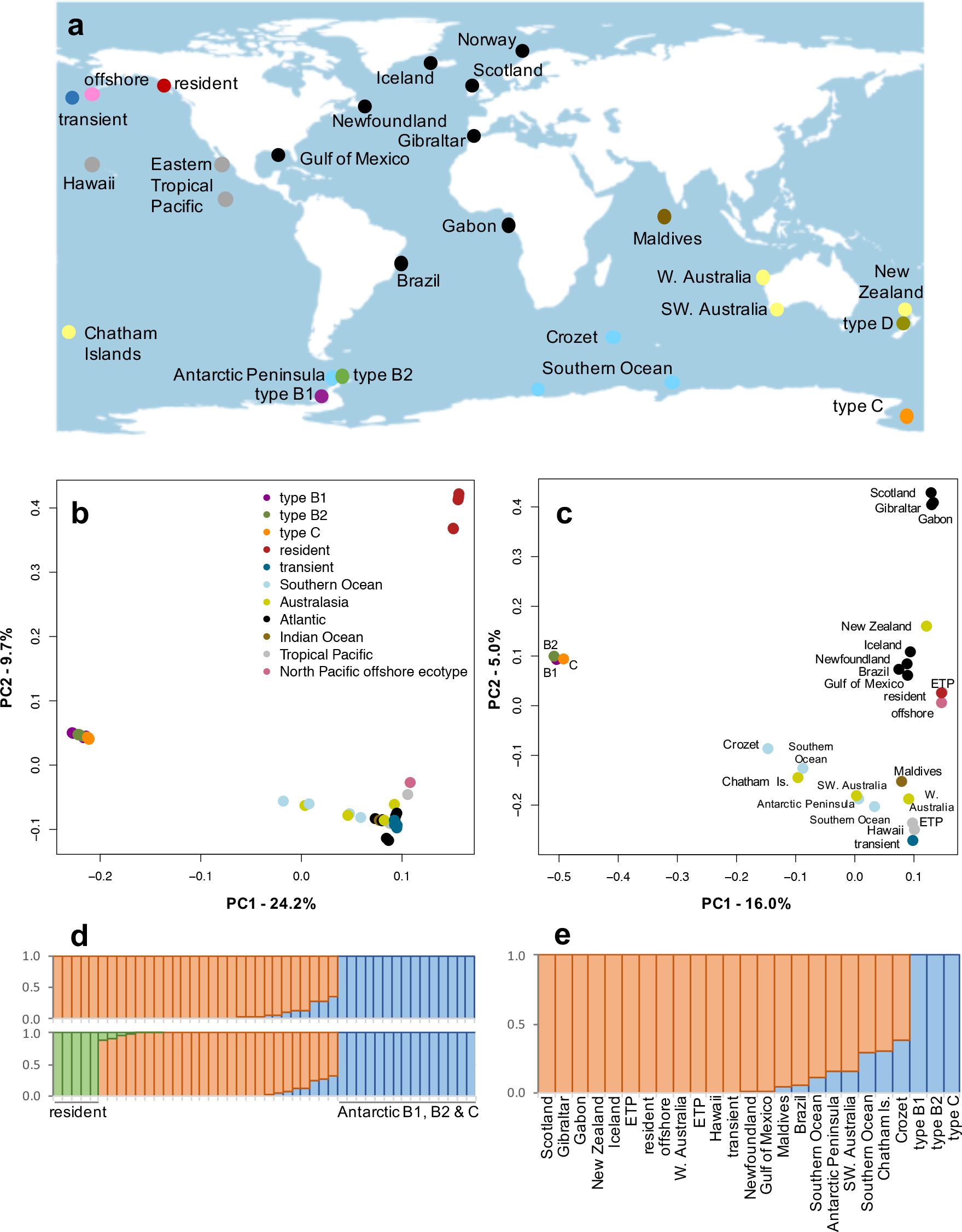
(a) Sampling locations of the individuals for which twenty-six 5× coverage genomes and one 27Mb partial genome were generated (global dataset). Marker colours are as per the PCA legend. An additional twenty low coverage genomes (ecotype dataset) were used in some analyses, see Foote et al. (2016) for sampling locations. (b) PCA plots of the combined global and ecotype datasets (excluding *type D*), and (c) when only one 5× coverage genome per population in included. (d) Individual admixture proportions, conditional on the number of genetic clusters (*K*=2 and *K*=3), for the combined global and ecotype datasets, and for (*K*=2) (e) when only one 5× coverage genome per population is included.

### 2.2 Library building, sequencing and mapping

Samples were selected from a global dataset of 452 individuals that best represented the known global geographic and genetic diversity of this species (**Morin et al., 2015**). Where possible, we selected identifiable individuals from longitudinally studied populations, *e.g*. Crozet Archipelago (**Guinet & Tixier, 2011**); Gibraltar (**Esteban et al., 2016**); and Iceland (**Samarra & Foote, 2015**). DNA was extracted from skin biopsies, with the exception of the recently described *type D* morphotype (**Pitman et al., 2011**), from which dry soft tissue and powdered tooth were sampled from a specimen (#1077) which stranded on Paraparaumu Beach, New Zealand in 1955, and is now part of the collections of the Museum of New Zealand, Te Papa Tongarewa, Wellington. This is the only available sample for DNA analysis from this rarely observed morphotype, which has a pelagic circumpolar subantarctic distribution, making it logistically difficult to biopsy sample (**Pitman et al., 2011**; **Foote et al., 2013**; **Tixier et al., 2016**).

DNA was extracted using a variety of common extraction methods as per **Morin et al. (2015)**. Genomic DNA was then sheared to an average size of ~500 bp using a Diagenode Bioruptor Pico sonication device. The sheared DNA extracts were converted to blunt-end Illumina sequencing libraries using New England Biolabs (Ipswich, MA, USA) NEBNext library kit E6040L. Libraries were subsequently index-amplified for 20 cycles using a KAPA HiFi HotStart PCR kit (Kapa Biosystems, Wilmington, Ma. USA) in 50-μl reactions following the manufacturer’s guidelines. The amplified libraries were then purified using a QIAquick PCR purification kit (Qiagen, Hilden, Germany) and size-selected on a 2% agarose gel in the range 422-580 bp using a BluePippin instrument (Sage Science, Beverly, MA. USA). The DNA concentration of the libraries was measured using a 2100 Bioanalyzer (Agilent Technologies, CA, USA); these were then equimolarly pooled and sequenced across four lanes of an Illumina HiSeq4000 platform using paired-read PE150 chemistry and two lanes using single-read SR100 chemistry.

DNA libraries from the soft tissue and powdered bone of the *type D* museum specimen were initially pooled and shotgun sequenced across one lane of an Illumina HiSeq2000. This indicated a low endogenous content of DNA (1.5% in soft tissue libraries and 4.0% in powdered tooth libraries). Libraries were therefore subjected to two rounds of whole genome enrichment capture using genome-wide biotinylated RNA baits built from modern DNA by Arbor Biosciences, Ann Arbor, MI (**Enk et al., 2014**), prior to pooling and sequencing across a second lane of an Illumina HiSeq2000. This enriched endogenous content 24-fold, so that post-capture 30.8-96.5% of reads mapped to the reference genome (**Foote et al., 2015**).

However, this resulted in a high percentage of duplicate reads (77%) suggesting the libraries contained limited starting template of endogenous DNA (**Ávila-Arcos et al., 2015**). Thus, the consensus sequence was reduced to 0.02× mean depth of coverage after the removal of duplicate reads. In total, 27,162,031 sites were covered at ≥1×. Analyses of a 41Mb scaffold (KB316842.1) using mapDamage2 (**Jónsson et al., 2013**) revealed that sequencing reads also exhibited characteristic post-mortem damage patterns (**Supporting Information Figure S1**). Specifically, an excess of C>T transitions at the 5’ termini as expected from deamination, and the complementary G>A transitions at the 3’ termini. Therefore, all downstream analyses on the *type D* sequence were restricted to transversions.

Read trimming, mapping, filtering and repeat-masking was conducted as per **Foote et al. (2016)**, with the exception that the previous study masked reads with a combined coverage of 200×, here we masked regions of low (less than a third of the mean) and excessive (more than twice the mean) combined coverage; regions of poor mapping quality (Q<30); and regions called as Ns in the reference sequence, all assessed using the CALLABLELOCI tool in GATK (**McKenna et al., 2010**; **DePristo et al., 2011**) and subsequently masked using BEDtools (**Quinlan & Hall, 2010**). Sites were further filtered to include only autosomal regions, except where stated otherwise, and only sites with a base quality scores >30 were used in all downstream analyses.

Changes in the cluster generation chemistry from bridge amplification to exclusion amplification has been reported to result in increased leakage of reads between indexes in pooled samples on a lane of the new Illumina sequencing platforms, which include the HiSeq4000 (**Sinha et al., 2017**). To assess the extent of leakage between indexes we mapped the sequencing reads from two HiSeq4000 lanes to a haploid killer whale mitochondrial genome (KF164610.1). Mitochondrial genomes had previously been sequenced using the 454 Life Sciences (Roche) and Illumina HiSeq2000 sequencing platforms (**Morin et al., 2010**; **2015**) and were used as a reference panel. The consensus sequence generated from the HiSeq4000 sequencing reads for this study, were 100% identical for the same individuals in the reference panel (**Morin et al., 2010**; **2015**). We then quantified contamination from leaked reads based on the protocol for assessing the extent of human contamination in Neanderthal sequenced data (**Green et al., 2010**). We inspected reads from a North Pacific ‘*offshore*’ killer whale, which mapped to the mitochondrial genome at 695× mean coverage (±72 S.E.) after filtering to remove low-quality bases (Q<30). At sites where the nucleotide was known to be private to the *offshore* sequence, we checked the counts of reads that concurred with the 454 Life Sciences (Roche) sequence generated from long-range PCR amplicons for the same individual (**Morin et al., 2010**), and the counts of reads that matched sequences of one or more of the other individuals pooled on the same HiSeq4000 lane. Counts of the mismatch alleles were uniformly low (mean < 0.5% of reads per site). Our results therefore concur with recent studies (**Owens et al., 2018**; **van der Valk et al., 2018**) that the rate of index swapping is low on the new Illumina platforms, and provide confidence that leakage between indexes did not greatly influence the inferred genotype likelihoods or the inferred genetic relationship among individuals.

### 2.3 Principal component analysis

The relationship of the samples in the global dataset to the killer whale ecotypes were explored using PCAngsd, a Principal Component Analysis (PCA) for low depth next-generation sequencing data using genotype likelihoods (GLs), thereby accounting for the uncertainty in the called genotypes that is inherently present in low-depth sequencing data (**Meisner & Albrechtsen, 2018**). We restricted the analyses to autosomal scaffolds ≥10Mb in length, which accounted for 1.5 Gb (~64%) of the genome (prior to masking). First, we estimated covariance of the 46 modern samples, which included five samples each for *resident*, *transient*, types *B1*, *B2* and *C*. After pruning SNPs to reduce linkage, a total of 225,281 SNPs were considered in this analysis. The eigenvectors from the covariance matrix were generated with the *R* function ‘eigen’, and significance was determined with a Tracy-Widom test (**Tracy & Widom, 1994**; **Patterson, Price & Reich, 2006**) performed in the R-package AssocTest (**Wang et al., 2017**) to evaluate the statistical significance of each principal component identified by the PCA. To reduce the influence of variable sample sizes among populations, we then repeated the analyses with a subset of 25 samples, removing four samples each from the *resident*, *transient*, types *B1*, *B2* and *C*, and removing the Norwegian sample, which belongs to the same metapopulation as the Icelandic sample (**Foote et al., 2011**).

To investigate the relationship between the *type D* morph and the other killer whale populations, a further PCA was constructed using the single read sampling approach implemented in ANGSD (**Korneliussen, Albrechtsen & Nielsen, 2014**). A random base was sampled from each position at which all samples were covered at ≥1× coverage to remove bias caused by differences in sequencing depth. Transitions were then removed due to the confounding effect of DNA damage patterns caused by deamination appearing as C→T transitions, and the corresponding reverse complement A→G in the sequence data. Random bases were therefore sampled from a total of 6,565 transversions.

### 2.4 Individual assignment and admixture analyses

An individual-based assignment test was performed using NGSadmix (**Skotte, Korneliussen & Albrechtsen, 2013**), a maximum likelihood method that bases its inference on genotype likelihoods. As for the PCA analysis, we ran the NGSadmix twice, once with the 46 modern samples (*i.e*. excluding the *type D* museum specimen) and once with the subset of 25 samples. As above, analyses were restricted to autosomal scaffolds ≥10Mb in length. NGSadmix was run with the number of ancestral populations *K* set from 1-10. For each of these *K* values, NGSadmix was re-run five times for each value of *K*, and with different seeds to ensure convergence. SNPs were further filtered to include only those covered in at least 25 individuals with a probability of *P* < 0.000001 of being variable as inferred by the likelihood ratio test and removing sites with a minor allele frequency of 0.05, so that singletons were not considered. Finally, SNPs were pruned to account for linkage, resulting in the analyses being based on 290,309 variant sites.

### 2.5 Distance-based phylogenetic inference

The genetic relationships among individuals within the dataset were further reconstructed with ngsDist (**Vieira et al., 2015**) using distance-based phylogenetic inference based on pairwise genetic distances. ngsDist takes genotype uncertainty into account by avoiding genotype calling and instead uses genotype posterior probabilities estimated by ANGSD. A block-bootstrapping procedure was used to generate 100 distance matrices, obtained by repetitively sampling blocks of 500 SNPs from the original data set of 6,974,134 SNPs. Pairwise genetic distances among the 5× coverage genomes were visualised as a phylogenetic tree using the distance-based phylogeny inference program FastME 2.1.4 (**LeFort, Desper & Gascuel, 2015**).

### 2.6 Pairwise sequentially Markovian coalescent

We used seqtk (https://github.com/lh3/seqtk) to combine 32 haploid male X-chromosome scaffolds of >1Mb each and totalling 91Mb, to construct pseudo-diploid sequences. The PSMC model estimates the Time to Most Recent Common Ancestor (TMRCA) of segmental blocks of the genome and uses information from the rates of the coalescent events to infer *N*_e_ at a given time, thereby providing a direct estimate of the past demographic changes of a population (**Li & Durbin, 2011**). The method has been validated by its successful reconstructions of demographic histories using simulated data and genome sequences from modern human populations (**Li & Durbin, 2011**). A consensus sequence of each bam file was then generated in fastq format sequentially using the SAMtools mpileup command with the -C50 option to reduce the effect of reads with excessive mismatches (**Li et al., 2009**); bcftools view -c to call variants; lastly, vcfutils.pl vcf2fq to convert the vcf file of called variants to fastq format. Pairs of fastq files were then merged using seqtk and PSMC inference carried out using the recommended input parameters for human autosomal data (**Li & Durbin, 2011**), i.e. 25 iterations, with maximum TMRCA (*T*max) = 15, number of atomic time intervals(*n*) = 64 (following the pattern 1*4 + 25*2 + 1*4 + 1*6), and initial theta ratio (*r*) =5. Plots were scaled to real time as per (**Li & Durbin, 2011**), assuming a generation time of 25.7 years (**Taylor et al., 2007**) and a neutral mutation rate of the X-chromosome (*μ*X) derived as *μ*X=*μ*A[2(2+α)]/[3(1+α)], assuming a ratio of male-to-female mutation rate of α = 2 (**Miyata et al., 1987**) and an autosomal mutation rate (*μ*A) of 2.34×10^−8^ substitutions/nucleotide/generation (**Dornburg et al., 2011**). This gave us an estimated *μ*X = 2.08×10^−8^ substitutions/nucleotide/generation. Only males were used in these analyses, which included *transient*, *resident*, Antarctic types *B1* and *C* as our focal ecotypes (our 5× coverage *type B2* genome sequence was generated from a female); and from our global dataset we included the sequences of samples from Gabon, Gibraltar, New Zealand, North Pacific *offshore* ecotype, Eastern Tropical Pacific (ETP)-Clipperton Island, Iceland, Gulf of Mexico, Brazil, Southern Ocean, SW. Australia, Chatham Islands, Crozet Archipelago, Hawaii, ETP-Mexico, and W. Australia.

### 2.7 Inferring admixture from D- and F-statistics

To investigate whether ecotype pairs evenly shared derived alleles with outgroup populations, or whether one ecotype shared an excess of derived alleles with outgroups suggesting either recent shared ancestry or introgression, we estimated the *D*-statistic (**Green et al., 2010**) for various combinations of ecotypes and outgroups. For example, if the sympatric North Pacific *resident* and *transient* ecotypes are considered to be the in-group, and *X* is a global outgroup sample, the test can be used to evaluate if the data are inconsistent with the null hypothesis that the tree (((resident, transient), X), dolphin) is correct and that there has been no gene flow between *X* and either *resident* or *transient*, or any populations related to them. The definition used here is from **Durand et al. (2011**):

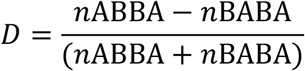

Where in the tree given above, *n*ABBA is the number of sites where *resident* shares the ancestral allele with the dolphin, and *transient* and *X* share a derived allele (ABBA sites); and, *n*BABA is the number of sites where *transient* shares the ancestral allele with the dolphin, and *resident* and *X* share a derived allele (BABA sites). Under the null hypothesis that the given topology is the true topology, we expect an approximately equal proportion of ABBA and BABA sites and thus *D* = 0. Hence a test statistic that differs significantly from 0 provides evidence either of gene flow, or that the tree is incorrect. The significance of the deviation from 0 was assessed using a *Z*-score based on jackknife estimates of the standard error of the D-statistics. This *Z*-score is based on the assumption that the D-statistic (under the null hypothesis) is normally distributed with mean 0 and a standard error achieved using the jackknife procedure. The tests were implemented in ANGSD and performed by sampling a single base at each position of the genome to remove bias caused by differences in sequencing depth at any genomic position. An error in our script reversed the sign of the value of D in a previous study (**Foote et al., 2016**), thus results differ in the direction of gene flow between that study and this, but do not change the conclusions drawn in that study, *i.e*. that some ecotypes are admixed.

The *f*_3_-statistic is based on the quantification of genetic drift (change of allele frequencies) between pairs of populations in a tree using variance in allele frequencies (**Reich et al., 2009**; **Patterson et al., 2012**; **Peter, 2016**). The *f*_3_-statistic can provide evidence of admixture, even if gene flow events occurred hundreds of generations ago (**Patterson et al. 2012**). These tests are of the form *f*_3_(A;B,C), where a significantly negative value of the *f*_3_ statistic implies that population A is admixed (**Patterson et al., 2012**). *f*_3_-statistics were computed using the estimators described in **Patterson et al. (2012)**, obtaining standard errors using a block jack-knife procedure over blocks of 1,000 SNPs.

When a potentially admixed population is identified the admixture proportions can be estimated using the ratio of *f*_4_-statistics. The expected value of the statistic *f*_4_(A,B; C,D) would be zero if we see no overlap in the paths of allele frequency changes between A and B, and between C and D through the tree. The expected value of the statistic *f*_4_(A,B; C,D) will be negative and significantly different from zero if allele frequency changes between A and B and between C and D take paths in the opposite direction along a shared edge within the tree; or positive and significantly different from zero if the drift between A and B and between C and D share overlapping paths in the same direction along an edge within the tree. The *f*_4_-statistic is not sensitive to post-admixture drift and can provide evidence of admixture, even if gene flow events occurred hundreds of generations ago (**Patterson et al., 2012**). To better identify the source populations that have admixed with the North Pacific *transient* ecotype we estimated *f*_4_(NZ, pilot whale; *X*, *transient*) and compared these with an estimate of *f*_4_(NZ, pilot whale; *resident*, *transient*). **Patterson et al. (2012)** defined the *f*_4_-ratio test as:

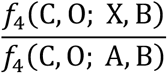

Where A and C are a sister group, B is sister to (A,C), X is a mixture of A and B, and O is the outgroup. This ratio estimates the ancestry from A, denoted as α, and the ancestry from B, as 1-α. It important to remember that neither the *transient* nor *resident* represent unadmixed lineages (see results), however, they do appear from past studies to represent the two most differentiated populations in the North Pacific (**Hoelzel et al., 2007**; **Parsons et al., 2013**; **Morin et al., 2010, 2015**). *f*_4_-statistics were computed using the estimators described in **Patterson et al. (2012)**, obtaining standard errors using a block jack-knife procedure over blocks of 1,000 SNPs.

### 2.8 Detecting archaic tracts

The prevalence of private alleles in Antarctic types *B1*, *B2* and *C*, as identified by the PCA, suggested potential archaic introgression from a now extinct (or otherwise unsampled) killer whale lineage or sister taxon. Archaic tracts with a distinctly older TMRCA than the genome-wide average can be identified even without an archaic reference genome. Private alleles resulting from *de novo* mutation along the branch to the Antarctic should be approximately randomly distributed across the genome, whereas tracts introgressed from a divergent lineage after vicariance of the Antarctic types from other populations, or differentially sorted from structure in an ancestral population will contain clusters of private alleles, the density of which will depend upon the divergence time of the introgressing and receiving lineages (**Racimo et al., 2015**). Since tract length is a function of recombination rate and time, tracts from ancestral structure are expected to be shorter than recently introgressed tracts due to the action of recombination (**Racimo et al., 2015**).

We therefore set out to screen for genomic tracts of consecutive or clustered private alleles in the Antarctic types. To ensure the results are comparable despite variation between samples in coverage at some sites, we randomly sampled a single allele at each site from each diploid modern genome. For the outgroup we used all variants found in a dataset consisting of the following widely distributed non-Antarctic samples: Gabon, Gibraltar, New Zealand, North Pacific *offshore*, *resident* and *transient* ecotypes, ETP-Clipperton Island, Iceland, Norway, Newfoundland, Hawaii, ETP-Mexico, Scotland and W. Australia; *i.e*. samples that show no evidence of admixture with the Antarctic types in either the PCA or NGSadmix analyses. For the ingroup we used *type B2*, which from the *f*_3_-statistics appeared to be the least admixed of the Antarctic types. Thus, we consider only variants found in *type B2*, which are not found in the widely distributed 14 non-Antarctic samples listed above.

We then used a Hidden Markov Model (HMM) to classify 1 kb windows into ‘non-archaic’ and ‘archaic’ states based on the density of private alleles (**Skov et al., 2018**). The background mutation rate was estimated in windows of 100 kb, using the variant density of all variants in non-Antarctic populations. We then weighted each 1 kb window by the proportion of sites not masked by our RepeatMasker and CallableLoci bed files. The HMM was trained using a set of starting parameters based on those used for humans (**Skov et al., 2018).** We trained the model across five independent runs, varying the starting parameters each time to ensure consistency of the final parameter input. Posterior decoding then determines whether consecutive 1 kb windows change or retain state (‘archaic’ or ‘non-archaic’) dependent upon the posterior probability.

## 3 RESULTS

Our results highlight that the distinctiveness of the killer whale ecotypes as described in the Introduction section, reflects their demographic and evolutionary histories, which include deep ancestral splits masked by more recent admixture. The latter confounding the inference of the relationships among these populations as a simple bifurcating tree-like model.

### 3.1 Genome sequences

Short-read sequence data were generated for 27 individuals, resulting in a mean coverage of 5× coverage of the autosomal region of the killer whale genome for the 26 modern samples, and 27Mb covered at ≥1× from a 62-year old museum specimen of the *type D* subantarctic morph. For some analyses, these data were combined with 20 previously sequenced 2× coverage genomes from the North Pacific *transient* and *resident* ecotypes, and Antarctic types *B1*, *B2* and *C* (**Foote et al., 2016**) and ~20× coverage RAD-seq data (**Moura et al., 2015**).

### 3.2 Principal Component Analysis: genetic variation segregates in Antarctic types and resident ecotype

In a PCA that included five samples per ecotype, the Antarctic types (*B1*, *B2* and *C*) separated out from all other killer whales along PC1 (**Figure 1b**), which explained 24.2% of the variation (**Supporting Information Figure S2a**). The *resident* ecotype formed a distinct cluster which separated out from a third cluster containing the *transient* ecotype and all other samples along PC2 (**Figure 1b**), explaining 9.7% of the variation (**Supporting Information Figure S2a**). Both first and second components were statistically significant: *P* < 0.001. This result was replicated when published RAD-seq data for a sub-Antarctic Marion Island sample were included (**Supporting Information Figure S3a**). The *transient* ecotype partially segregates from other samples along PC4, which explains 2.6% of the total genetic variation (**Supporting Information Figure S4**).

Uneven sampling of different demes can influence the inference of population clusters in admixture and PCA plots (**McVean, 2009**; **Gilbert, 2016**; **Lawson, van Dorp & Falush, 2018**). For example, when the closely related Norwegian and Icelandic samples are both included in the PCA, they segregate from the other samples along PC2 (**Supporting Information Figure S5**). After reducing the dataset used in the PCA to one sample per population to reduce this bias, differences between the Antarctic types and all other killer whales continue to explain the greatest and only significant (*P* < 0.001) component of variation in the data (**Figure 1c**; **Supporting Information Figure S2b**). This pattern remains even when just a single *type B1* sample (*i.e*. no *B2* or *C* samples) is included, and likewise for single *B2* and *C* samples, albeit with less variation (~11%) explained by PC1 (**Supporting Information Figure S6**). Reducing the dataset to one sample per population results in a change in clustering along PC2, along which samples are distributed, to some extent, reflecting a cline from the North Atlantic to the North Pacific, but with the *resident* and *offshore* samples intermediate between Atlantic and Pacific samples (**Figure 1c**). A further PCA identified the *type D* morphotype (**Pitman et al., 2010**) as being genetically intermediate between Southern Ocean and North Atlantic populations (**Supporting Information Figure S7**).

### 3.3 Individual assignment and admixture analyses support PCA inference

The results of the PCAs are reflected in the admixture plots inferred by NGSadmix (**Figure 1d,e**). The uppermost hierarchical level of structure, inferred from the greatest step-wise increase in log likelihood (Δ*K*; **Evanno, Regnaut, & Goudet, 2005**) identified two clusters (**Supporting Information Figure S8**). As in the PCA, one cluster consisted of Antarctic types *B1*, *B2* and *C*, the other a mostly homogenous cluster of all other killer whales, albeit with ‘Antarctic’ ancestry detected in some southern hemisphere samples (**Figure 1d,e**). When multiple samples per ecotype are included, we find the second highest Δ*K* from *K*=2 to *K*=3 clusters, in which the North Pacific *resident* ecotype forms a discrete cluster (**Figure 1d**).

PCA and STRUCTURE-like admixture models use similar information and generate similar axes of variation (**Patterson, Price & Reich, 2006**; **Lawson, van Dorp & Falush, 2018**). Both methods are likely to identify the samples with the greatest population-specific drift that therefore share rare derived alleles or have lost ancestral alleles from standing variation, as the major axes of structure (**Lawson et al., 2018**). Accordingly, both PCangsd and NGSadmix identified the uppermost hierarchical level of structure within our dataset as being between the Antarctic types and all other killer whales (**Figure 1**). The spatial distribution of samples within the PCA plot can be considered as being representative of the mean pairwise coalescent times between each pair of samples (**McVean, 2009**). Changes in frequency or the loss of neutral alleles through population specific drift will result in more recent mean coalescence among individuals, thereby causing them to cluster together and segregate from other populations along the axes of the PCA (**McVean, 2009**). Our results are therefore consistent with previous findings of a shared demographic history of the Antarctic types that included a shared population bottleneck and substantial drift (**Morin et al., 2015**; **Foote et al., 2016**). However, our finding that this pattern in the PCA is retained when just a single Antarctic sample is included (**Figure S7**), indicates this pattern is not just driven by the shared loss of standing variation in the Antarctic types, but that alleles explaining a significant proportion of the observed genetic variation coalesce within that single sample in those analyses. This suggests that there is are a large number of alleles private to the Antarctic types contributing towards the pattern of global genetic variation in killer whales. It should be noted that at higher values of *K* further structuring is identified within our dataset; for example, at *K*=7 Antarctic types *B1*, *B2* and *C* form three discrete clusters as per **Foote et al.(2016)**. However, due to our sampling scheme, our focus in this study was not to identify regional structuring, but to identify the major axes of global structure and to infer the underlying processes.

### 3.4 ‘Archaic’ tracts in Antarctic types suggest ancient admixture with an ancient ‘ghost’ population

The HMM method for detecting archaic tracts based on private allele density (**Skov et al., 2018**) inferred 1,897 tracts totalling 8,119 kb of the tested 41 Mb scaffold as archaic in Antarctic *type B2* with a posterior probability of ≥0.5 (**Supporting Information Figure S9**). Thus, up to 21.6 % of the genomic region analysed was inferred to be potentially comprised of archaic ancestry. However, a proportion of these windows inferred to be in the archaic state with a posterior probability of ≥0.5 may be false positives. Considering windows inferred as archaic with posterior probabilities of ≥0.8 identified 18 archaic tracts totalling 224 kb.

The emission probabilities of the HMM are modelled as Poisson distributions with means of *λ_Archaic_* = *μ* · *L* ∙ *T_Archaic_* for the archaic state and *λ_Ingroup_* = *μ* · *L* · *T_Ingroup_* for the non-archaic (or ingroup) state (**Skov et al., 2018**), where *L* is the window size (1000 bp) and *μ* is the mutation rate (2.38 ×10^−8^; **Dornburg et al., 2011**). This allows us to estimate the mean TMRCA of the archaic and ingroup windows with the corresponding segments in the outgroup dataset. The TMRCA between the archaic tracts within *type B2* and the corresponding genomic regions in the outgroup is a Poisson distribution around a mean of 9,786 generations (~251 KYA, assuming a generation time of 25.7 years; **Taylor et al., 2007**). The estimated TMRCA between the non-archaic tracts within *type B2* and the corresponding genomic regions in the outgroup is a Poisson distribution around a mean of 2,429 generations (~62 KYA). Thus, the genome of *type B2* appears to be admixed, comprising of approximately 80% ancestry that coalesces with the ancestry of the outgroup during the previous glacial period (Marine Isotope Stage 5), and approximately 20% ancestry derived from an unsampled lineage that coalesces with the ancestry of the outgroup during an earlier glacial cycle (Marine Isotope Stage 8).

Tracts inferred to be in the archaic state with a posterior probability of ≥0.5 were on average between 5 and 6 windows long, *i.e*., between 5 and 6 kb; an order of magnitude shorter than introgressed archaic tracts in non-African humans (**Skov et al., 2018**). Considering just the tracts called as archaic with posterior probabilities of ≥0.8, the average tract length was 12-13 kb. The estimation of the time of introgression from tract length is dependent upon recombination rate (*r*), which has not yet been estimated for killer whales. However, assuming a constant value of *r* approximated to the mean *r* estimated for the human genome of 1.2 × 10^−8^/bp (1.2 cM/Mb; **Dumont & Payseur, 2008**) places the age of tracts 5.5 kb long to approximately 14,000 generations ago, *i.e*. older than *T_Archaic_* (**Supporting Information Figure S10**). A recombination rate of ≥5.0 × 10^−8^/bp (5.0 cM/Mb) would be required for such short tract lengths to have introgressed during the last 2,500 generations, *i.e*. close to *T_Ingroup_* (**Supporting Information Figure S10**). Considering tracts of 12.5 kb length suggests a time of introgression between *T_Archaic_* and *T_Ingroup_* of approximately 7,000 generations (assuming *r* = 1.2 × 10^−8^/bp). Assuming a recombination rate for killer whales in the range of humans thus suggests a scenario different from the recent introgression from Neandertals into the lineage of non-African humans. Instead, the source of archaic ancestry tracts in *type B2* killer whales is better explained by ancestral population structure. This therefore requires a scenario in which these tracts were the minor component of the ancestry (*i.e*., the lineage that contributed less to the gene pool, see **Schumer et al., 2018**) of an admixed ancestral killer whale population between *T_Archaic_* and *T_Ingroup_*, and this ancestry was therefore already being broken up by recombination prior to *T_Ingroup_* (**Figure 2**).

**Figure 2.**
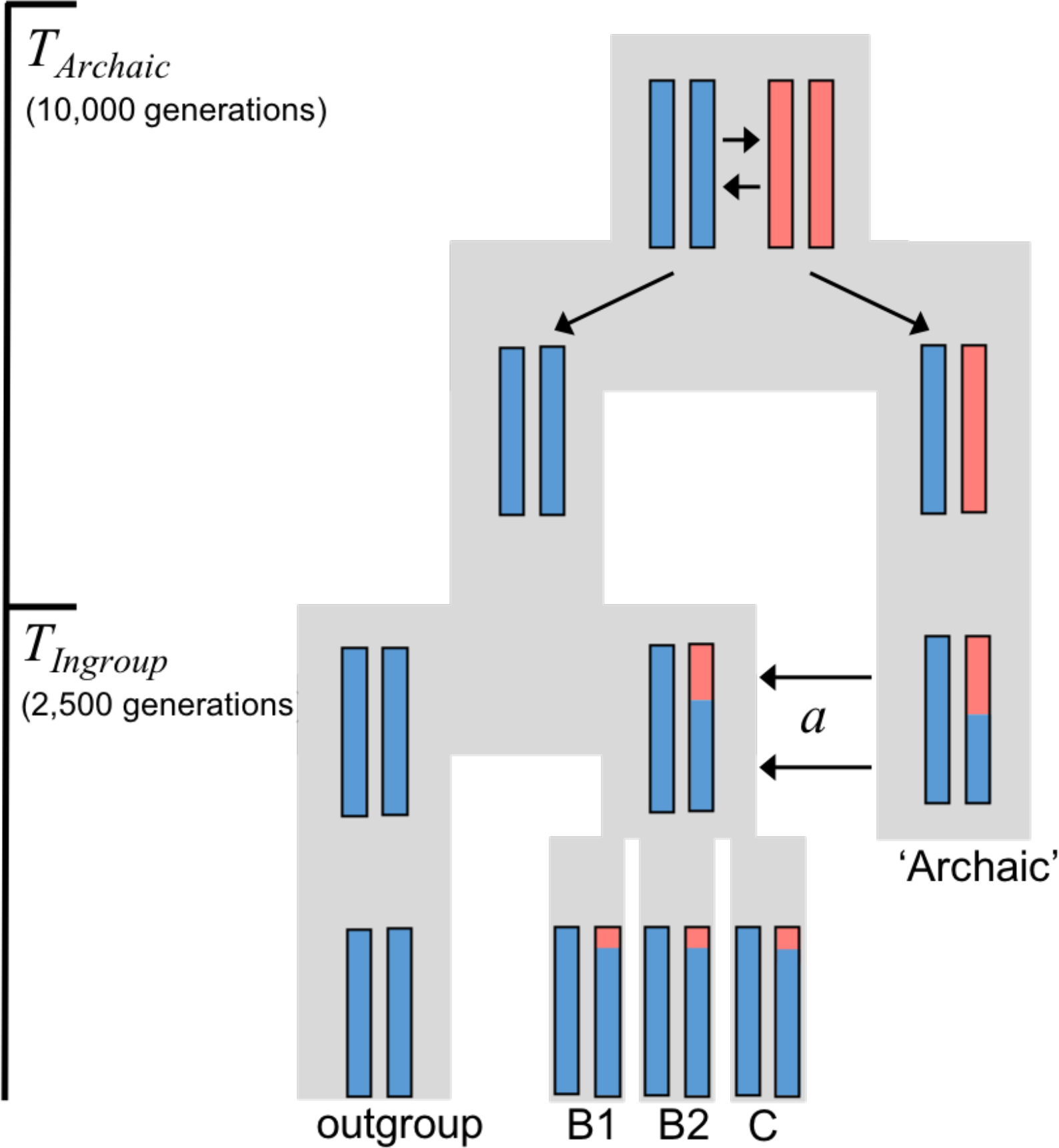
Model of hypothesised demographic scenario. Approximately 80% of the ancestry of *type B2* has a mean time to most recent common ancestor (TMRCA) of 2,500 generations with the outgroup samples (*T_Ingroup_*), whilst 20% of the ancestry of *type B2* is in short segments (5-6 kb) with a TMRCA of 10,000 generations with the outgroup samples (*T_Archaic_*). We therefore propose lineage sorting of ancestral structure subsequent to *T_Archaic_*, so that ancestry represented by red shading was not found in the most recent common ancestor of the outgroup and *type B2*. After *T_Ingroup_*, the red shaded ancestry introgressed into the ancestor of *type B2*, but not the outgroup, from an unsampled ‘archaic’ source lineage. Figure is adapted from Figure 1a of Skov et al. (2018) and Figure 1 of Racimo et al. (2015). Shading represents the decreasing length of the archaic (red) ancestry tracts through the action of recombination.

A PCA plot of SNPs occurring within the 1 kb windows inferred by the HMM as being in the archaic state (at posterior probability >0.5) highlighted the role of these archaic tracts in contributing to the major axis of structure in our global dataset (**Supporting Information Figure S11**). It also indicates variation among types *B1*, *B2* and *C* in the sharing of variants within these tracts. A PCA estimated from the non-archaic tracts (not shown) generated similar PCs and eigen values to **Figure 1c** and so the differentiation of the Antarctic types is not driven solely by the archaic tracts.

### 3.5 PSMC suggests an early split of *transients*, and ancestral vicariance and admixture in Antarctic types

To better understand the chronology of the divergence of killer whale ecotypes, we employed a method that drew inference from the distribution of the lengths of shared Identity-by-State (IBS) tracts to investigate coalescence rates through time. We created pseudo-diploid sequences by combining the phased haploid non-pseudoautosomal X-chromosome sequences from two different males and used PSMC (**Li & Durbin, 2011**) to estimate changes through time in the coalescent rate between the two X-chromosome haplotypes. The *y*-axis of a PSMC plot is driven by both changes in population structuring and demography and is more accurately interpreted as an estimate of the inverse of the rate of coalescence at any point in time represented along the *x*-axis (**Mazet et al., 2016**). Pseudo-diploids comprised of two haploid male X-chromosome sequences can therefore be used to infer the approximate population split time between two populations (**Li & Durbin, 2011**). When populations diverge and all gene flow between them ceases, the accumulation of new mutations and loss of diversity through drift will be population specific (**Figure 3a**). Population divergence therefore manifests itself in the pseudo-diploid sequence as heterozygote sites that break up long homozygous tracts from which more recent coalescent events are inferred. This results in an upsweep along the *y*-axis of the PSMC plot approximately at the point of cessation of gene flow (**Li & Durbin, 2011**). Post-divergence migration between the two demes being compared can result in a more recent coalescence of post-divergence mutations, shifting the upsweep closer to the present along the *x*-axis (**Cahill et al., 2016**).

**Figure 3.**
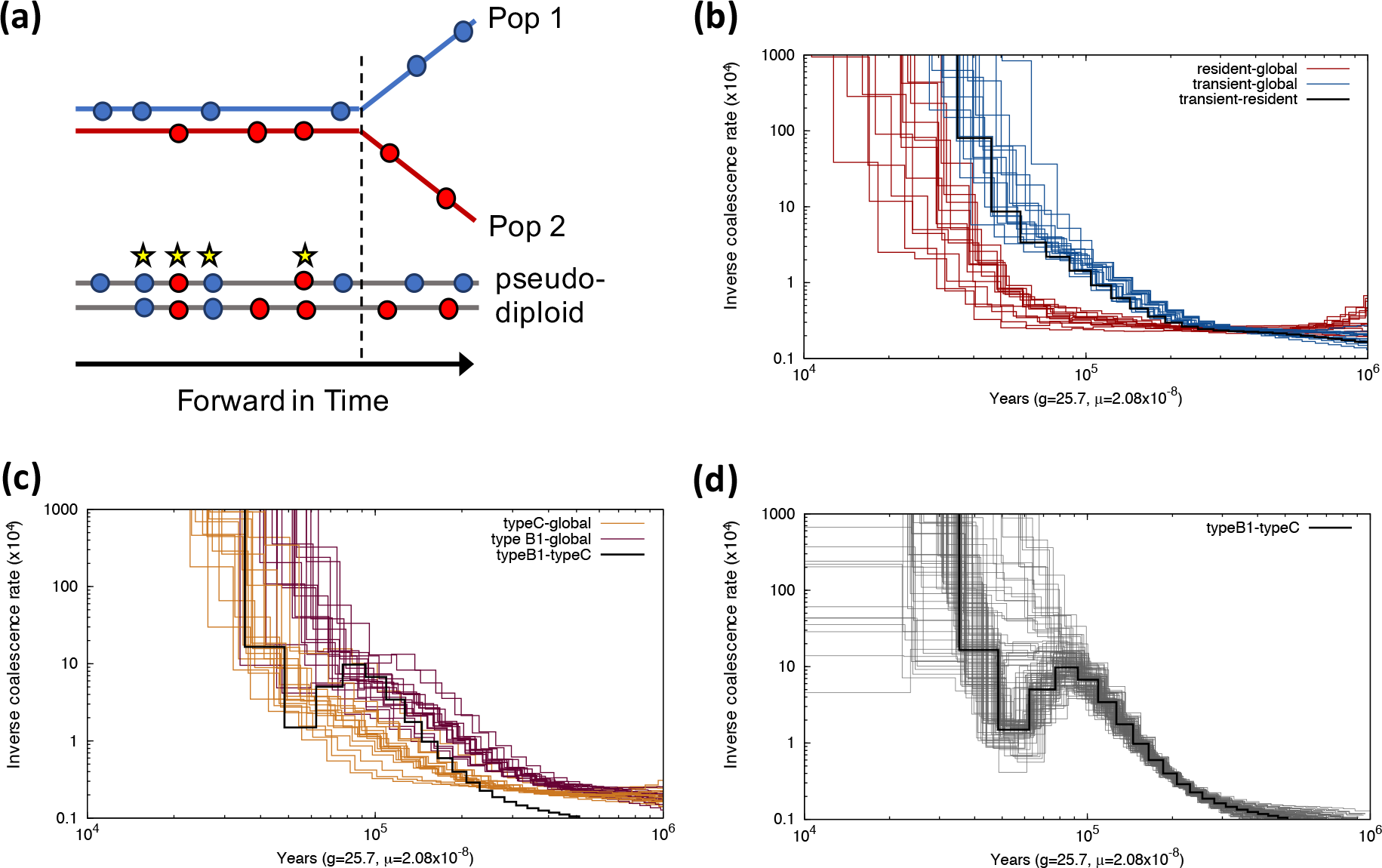
Pairwise sequentially Markovian coalescent (PSMC) plots of changes in coalescence rates between haploid male X-chromosomes combined to construct pseudo-diploid X-chromosomes. (a) A schematic diagram of the accumulations of mutations (indicated by circles) in two populations which are initially connected by gene flow, but diverge without further gene flow at the time indicated by the dashed line. A pseudo-diploid comprised of haploid chromosomes from Pop1 and Pop2 can be homozygous for the derived alleles at mutations when there is gene flow and recombination (indicated by yellow stars) between Pop1 and Pop2. Mutations accumulated after the cessation of gene flow will remain private to Pop1 or Pop2 and therefore inferred as heterozygotes in the pseudo-diploid. The accumulation and distribution of heterozygotes in the pseudo-diploid breaks up homozygous tracts which PSMC infers as a decrease in the coalescence rate. Therefore, the exponential upsweep towards infinity on the y-axis of the PSMC plot of a pseudo-diploid genome provides a coarse measure of divergence time. (b) The haploid X-chromosome of a male *resident* (red) and a male *transient* (blue) are combined with haploid X-chromosomes of males from the global dataset: Gabon, Gibraltar, New Zealand, North Pacific *offshore* ecotype, Eastern Tropical Pacific (ETP)-Clipperton Island, Iceland, Gulf of Mexico, Brazil, Southern Ocean, SW Australia, Chatham Islands, Crozet Archipelago, Hawaii, ETP-Mexico, and W. Australia. Each pseudo-diploid is represented by a separate plot. (c) PSMC plots of pseudo-diploid X-chromosome constructed from the haploid X-chromosome of either a male *type B1* (aubergine) or a male *type C* (orange) together with the haploid X-chromosome of a global sample (as for panel b). The plot of the combined B1-C pseudo-diploid X-chromosome is shown in black and shown separately in (d) with the thick black line representing the median and thin grey lines corresponding to 100 rounds of bootstrapping. Inverse coalescence rate is scaled by 2*μ*.

Applying this approach, we infer decreasing coalescence from the upsweep in estimated *N*_e_ from 200-300 KYA between the *transient* and all tested populations (**Figure 3b**). In contrast, coalescence does not appear to decrease between the *resident* and these same populations until approximately 100 KYA (**Figure 3b**). In other words, the *resident* shares a higher proportion of longer IBS tracts within the X-chromosome with the tested global samples, representing more recent recombination events, than the *transient* does with those same populations. Or to put it another way, the mean TMRCA of the X-chromosome is older between the *transient* ecotype and the populations tested here, than between the *resident* ecotype and those same populations. We interpret this as an earlier matrilineal fission and divergence from these globally distributed samples by the ancestor of the present-day *transient* ecotype, and a later founding of the ancestor of the present-day *resident* ecotype. This is consistent with earlier estimates of TMRCA based on mitochondrial genomes (**Morin et al., 2015**) and the inferred timing of founder bottlenecks based on nuclear genomes (**Foote et al., 2016**).

Comparing coalescent patterns of the X-chromosome of a *type B1* male and a *type C* male with the global dataset we find that the B1-global pseudo-diploid plots follow a similar trajectory to the transient-global plot, whereas the C-global pseudo-diploid plot upsweep suggests a more recent decrease in coalescence with the global samples (**Figure 3c**). Thus, despite the covariance of allele frequencies and resulting clustering in the PCA and admixture plots (**Figure 1**), *type B1* and *type C* differ in their sharing of shorter IBS segments with a TMRCA >100 KYA. An upsweep of inferred *N*_e_ in plot of the types *B1* and *C* pseudo-diploid commencing between 200-300 KYA, stalls at approximately 90 KYA and declines between 90-50 KYA, before increasing again (**Figure 3c**). The increase in coalescence, estimated between 50-90 KYA, implies a period of admixture between types *B1* and *C*. The bootstrap plots illustrate the variation in ancestry across the X-chromosome (**Figure 3d**).

### 3.6 Comparing mitochondrial DNA (mtDNA) and nuclear DNA (nuDNA) tree topologies

A distance-based tree estimated from pairwise genetic differences is only partially concordant with the mitochondrial DNA tree (**Supporting Information Figure S12**). Thus, in some cases genetic variation of the nuclear genome is shared among samples reflecting the matrilineal fission process that drives divergence in social and population structure in most killer whale populations studied to date (**Ford, 2009**); in other cases, geographically proximate samples with divergent mtDNA haplotypes cluster in the nuclear tree, suggesting a role for long-range matrilineal dispersal and subsequent gene flow in shaping patterns of nuclear genome diversity.

### 3.7 D-statistics indicate Antarctic types B1, B2 and C differ in their sharing of derived alleles with outgroup populations

The *D*-statistic (**Green et al., 2010**) considers a tree-like relationship among four populations, *e.g*. (*ecotype1*, *ecotype2*; *X*, dolphin), and estimates whether *X* shares an excess of derived alleles with one of the two ecotypes in the ingroup. Significant sharing of derived alleles between an in-group and *X* indicates either introgression from *X* (or a closely related population) into one ecotype, but not the other; or that the tree topology is incorrect, and *X* belongs in the in-group. Estimation of *D*(*type B2*, *type C*; *X*, dolphin) found that 19 out of 23 tests were considered significant based on *Z*-score > 3, and that these 19 samples shared an excess of derived alleles with *type C* relative to *type B2* (**Figure 4a**). A similar result was obtained when *type B2* was swapped for *type B1*, *i.e*. *D*(*type B1*, *type C*; *X*, dolphin); in this test *type C* shares a significant excess of derived alleles with all outgroups (*X*), except the sample from the Crozet Archipelago. There was no significant difference between *type B1* and *type B2* in the sharing of alleles derived in any of the outgroup samples (*X*).

**Figure 4.**
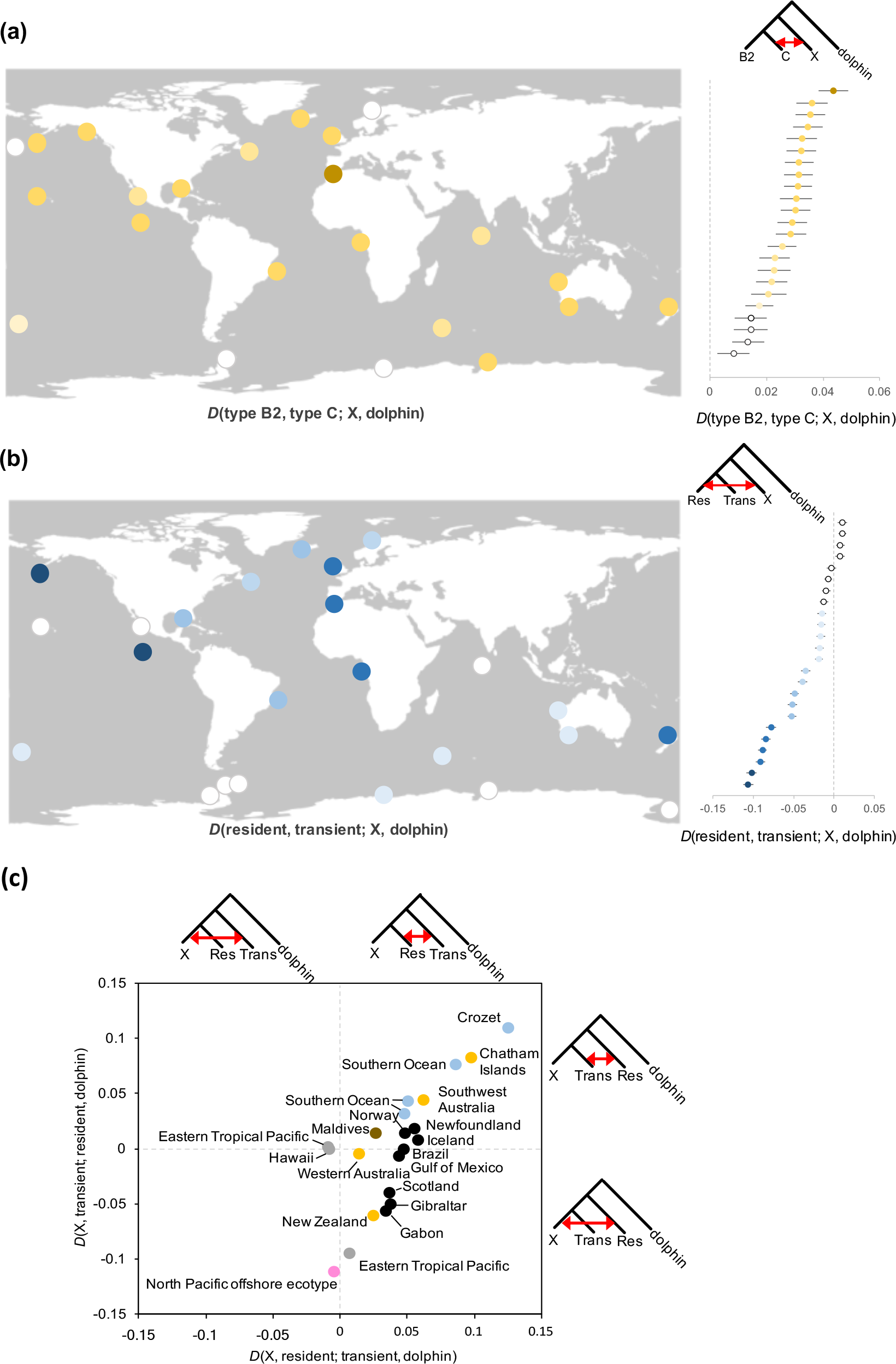
(a) Samples on the map are coloured by the value of D(type B2, type C; X, dolphin). The nineteen highest positive statistics were considered statistically significant following correction for multiple testing, based on Z-scores >3. This indicates that in these nineteen tests X shares an excess of derived alleles with Antarctic *type C* relative to *type B2*. Standard errors are shown as horizontal bars on the markers in the plot to the right. (b) Estimates of D(resident, transient; X, dolphin), in which the sixteen most negative statistics were considered statistically significant following correction for multiple testing, based on Z-scores < −3. This indicates that in these sixteen tests X shares an excess of derived alleles with the North Pacific *resident* ecotype relative to the North Pacific *transient* ecotype. Standard errors are shown as horizontal bars on the markers in the plot to the right. (c) A comparison of D(X, resident; transient, dolphin) and D(X, transient; resident, dolphin). Negative values along the x-axis indicate samples which shared an excess of derived alleles with the *transient* ecotype. Negative values along the y-axis indicate samples which shared an excess of derived alleles with the *resident* ecotype.

### 3.8 D-statistics indicate Pacific *resident* and *transient* ecotypes differ in their sharing of derived alleles with outgroup populations

Estimation of *D*(*transient*, *resident*; *X*, dolphin) found that while none of the global outgroup samples shared a significant excess of derived alleles with the *transient* ecotype, a widely geographically distributed set of 16 out of 24 samples shared a significant (*Z*-score < −3) excess of derived alleles with the *resident* ecotype (**Figure 4b**). This may indicate that the topology (*transient*, *resident*; *X*, dolphin) is incorrect, and that in these 16 tests (*resident, X*) is the correct in-group, which would be consistent with the hypothesis of secondary genetic contact between the *transient* and *resident* ecotypes (**Foote et al., 2011**). We therefore compared *D*(*X*, *resident*; *transient*, dolphin) and *D*(*X*, *transient; resident*, dolphin) to assess this possibility (**Figure 4c**).

When we consider alleles purportedly derived in the *transient* (*i.e*. where *X* and *resident* are the ingroup) there is no significant sharing of excess derived alleles between any population represented by (*X*) and the *transient* ecotype. However, when *X* was a non-North Pacific sample, there was significant sharing of derived alleles between the *resident* and *transient* ecotypes (**Figure 4c**). When we consider *X* and the *transient* as the ingroup (*i.e*. alleles purportedly derived in the *resident*) the North Pacific *offshore*, ETP-Clipperton Island, New Zealand, Gibraltar, Gabon and Scotland samples all shared a significant excess of derived alleles with the *resident* ecotype (**Figure 4c**). These same populations generated the most strongly negative D-statistics in the test *D*(*transient*, *resident*; *X*, dolphin) (**Figure 4b**) and share a more recent common maternal ancestor with the *resident* than the *transient* ecotype based on mitochondrial genome phylogeny (**Morin et al., 2015**; **Supporting Information Figure S12**). We therefore interpret these results as a further indication that the *resident* ecotype diverged more recently from these six populations than it did from the *transient*, but that gene-flow between the *resident* and *transient* has subsequently occurred, most likely within the North Pacific. A comparison of *D*(*transient*, *resident*; *X*, dolphin) and *D*(*transient*, *offshore*; *X*, dolphin) indicates correlated (Pearson’s correlation coefficient: *r*_23_ = 0.9609, *p* < 0.00001) sharing of derived alleles between the *resident* and *X*, and between the *offshore* and *X* (**Supporting Information Figure S13**) supports this inference of recent shared ancestry between the *resident* and *offshore* ecotypes.

### 3.9 F-statistics indicate admixture between *transient* and *resident* lineages

The *f*_3_-statistic is based on the quantification of covariance of allele frequencies (often referred to as shared drift) between pairs of populations in a tree using variance in allele frequencies (**Reich et al., 2009**; **Patterson et al., 2012**). To identify admixed ecotypes, we estimated *f*_3_-statistics of the form *f*_3_(ecotype; *X*, *Y*). Significantly negative *f*_3_-statistics indicate varying levels of admixture between the ecotype and *X*, and between the ecotype and *Y*, so the estimate of allele frequency differences between the ecotype and *X* are negatively correlated with the differences in allele frequencies between the ecotype and *Y*. *F*_3_-statistics were minimized and significantly (*Z*-score < −3) negative when estimating *f*_3_(*transient*; *X*, *Y*), with the exception of when both *X* and *Y* were Antarctic types (**Figure 5**, **Supporting Information Figure S14**). This result indicates the *transient* ecotype is admixed with one or more of the donor populations, or with closely related populations with partly shared derived ancestry with the donor population (see for example the outgroup case in **Patterson et al., 2012**). The most negative *f*_3_-statistics were estimated for tests when Hawaiian and/or Mexican ETP samples represented *X* and *Y* (see columns 1 & 2 of the lower diagonal of **Figure 5**), consistent with the results from PCA (**Figure 1c**) and D-statistics (**Figure 4b,c**). Based upon *f*_4_-ratio tests, the Hawaiian and Mexican ETP samples shared a higher proportion of *transient* than *resident* ancestry (**Supporting Information Figure S15**).

**Figure 5.**
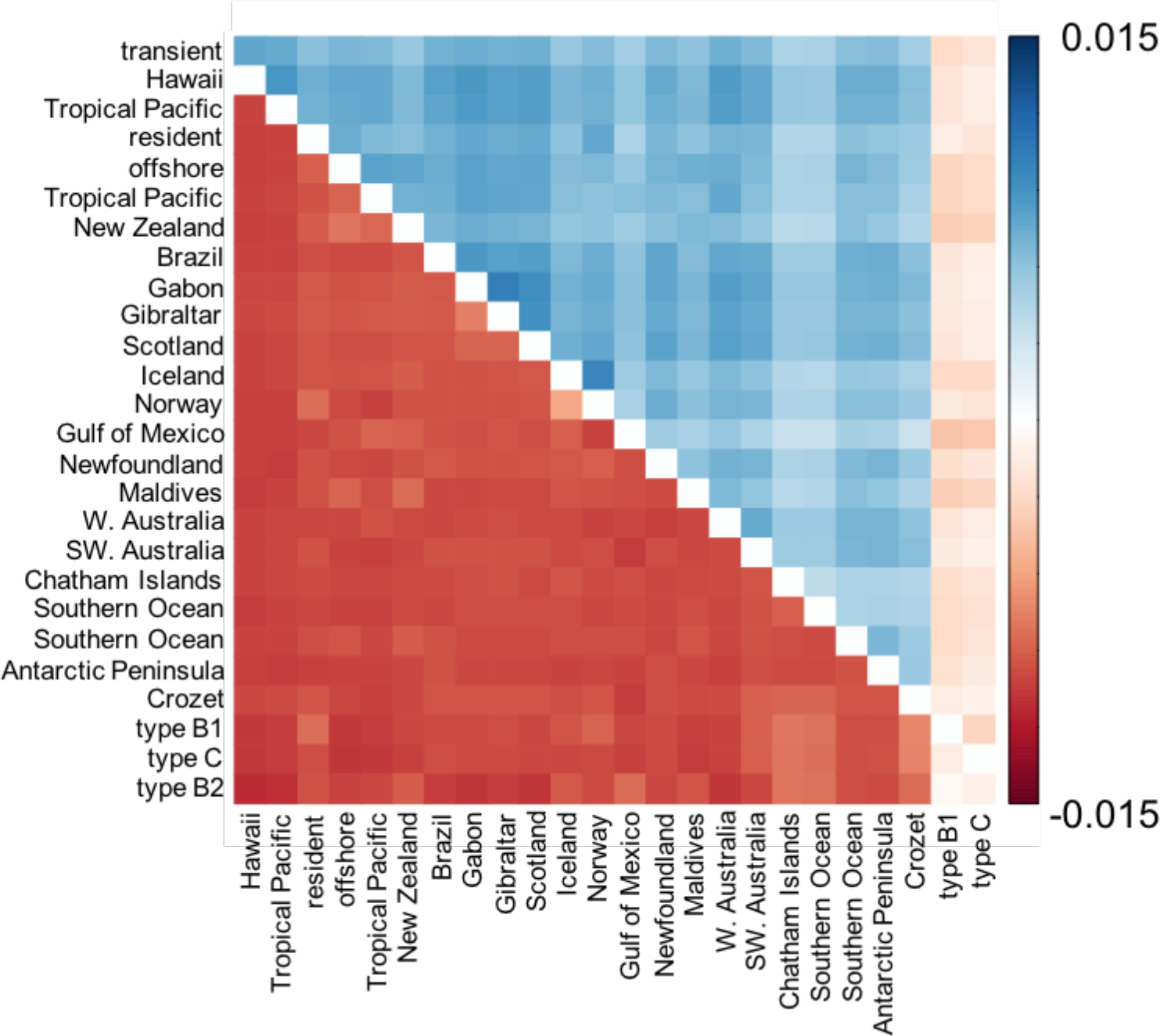
*F*_3_-statistics of the form *f*_3_(ecotype; *X*, *Y*), showing *f*_3_(*type B2*; *X*, *Y*) in the upper diagonal and *f*_3_(*transient*; *X*, *Y*) in the lower diagonal. The negative values for *f*_3_(*transient*; *X*, *Y*) indicate that the *transient* ecotype is highly admixed by X and/or Y, or population(s) closely related to them. The positive values for *f*_3_(*type B2*; *X*, *Y*) indicate that post-admixture drift from *X* and *Y* is greater than any admixture with *X* or *Y*, with the exception of *type B1* or *type C*.

### 3.9 F-statistics indicate drift is greater than any admixture in Antarctic types

*F*_3_-statistics were positively maximized when Antarctic *type B2* was the target ecotype for all tested combinations of *X* and *Y* (**Figure 5**, **Supporting Information Figure S14**). Positive and non-significant *f*_3_-statistics can arise despite admixture, for example, if population specific post-admixture drift in the target population is so large, it masks gene flow from the tested donor populations (**Patterson et al., 2012**). The extent of drift in *type B2* is such that it acts as an outgroup in *f*_3_-statistics, the same way that African genomes (*e.g*. Yoruban or Mbuti) are often used as an outgroup when comparing shared ancestry of Eurasian populations as *X* and *Y* in studies of human populations (*e.g*. **Seguin-Orlando et al., 2014**; **Pagani et al., 2016**). Therefore, we estimate the highest positive *f*_3_-statistics for tests when *X* and *Y* are known to originate from closely related populations, for example, *f*_3_(B2; Norway, Iceland) (**Figure 5**).

## 4 DISCUSSION

The present-day major axes of global genetic structure in killer whales are associated with the strongest drift having occurred in populations at the high latitude extremes of the species range. This pattern likely reflects some of the major demographic events in the last tens of millennia of the history of this species. Consistent with the expectations of range expansion theory (**Excoffier, Foll & Petit, 2009**) we find that populations that have expanded into areas inaccessible during the LGM (*e.g*. Antarctica and the Northern range limits of the Atlantic and Pacific) or have undergone some other long-range dispersal (*e.g*. the *resident* ecotype) have the greatest differentiation from neighbouring populations, indicating they have undergone the greatest drift in allele frequencies. The same high latitude populations show expansion from a small ancestral founder lineage based on the TMRCA of the mitochondrial genome (**Morin et al., 2015**) and coalescence patterns in the nuclear genome (**Foote et al., 2016**). Our results therefore expand the model of the evolution of population structure in the North Pacific proposed by **Hoelzel et al. (2007),** *i.e*. strong founder effects from ancestral colonising matrilines, to partially explain the major axes of structure in killer whales at high latitudes including Antarctica and the Northeast Atlantic (Norway and Iceland). However, despite these commonalities in the demographic and evolutionary histories of high latitude killer whale populations, the Antarctic types *B1*, *B2* and *C* stand out as explaining by far the most significant component of global genomic variation in this species.

We find an additional source of genetic variation in the Antarctic types in the form of private alleles clustered within short archaic tracts. The majority of the genome of the Antarctic types coalesces in the shared ancestral bottlenecked population and has a mean TMRCA of approximately 60 KYA with a widely distributed dataset of outgroup samples (**Figure 2**).

However, we also find short ancestry tracts which have an estimated mean TMRCA of over 200 KYA with corresponding genomic regions in these same outgroup samples (**Figure 2**). Thus, the genomes of Antarctic killer whales contain ancestry reflecting deeper coalescence during a previous glacial cycle. This is supported by our analysis of changes in coalescence rate through time between X-chromosome haplotypes of Antarctic types *B1* and *C* using PSMC, which although coarse, also indicate two peaks in coalescence, one at ~60 KYA and another >200 KYA. There is further support for an older and younger coalescence among Antarctic types from the TMRCA of mitochondrial genomes (see **Supporting Information Figure S16**; **Morin et al., 2015**). We hypothesize that this pattern in the mtDNA phylogeny is the result of replacement of mtDNA haplotype diversity of the Antarctic types during this episode of admixture between 50-90 KYA inferred from the PSMC plot (**Figure 2d**). The fixation of an introgressed mtDNA haplotype would be dependent upon admixture rate and effective population size (see **Posth et al., 2017**); high admixture rates and low *N*_e_ would be needed to drive the near fixation of the ancestral mtDNA lineage in types *B1*, *B2* and *C* and the pattern observed in the *B1*-*C* pseudo-diploid PSMC plot. Considering all these lines of investigation together, we interpret the results as being indicative of cyclical range expansions and contractions concurrent with the glacial cycles. Antarctic populations would be able to expand their range southwards during inter-glacial periods, increasing genetic differentiation from lower latitude populations, but then would retreat northwards during glacial periods, increasing contact and gene flow with lower latitude populations.

Our finding that the strongest structuring in a global dataset of killer whales is between the ecotypes found around Antarctica (types *B1*, *B2* and *C*) and all other killer whales counters claims by **de Bruyn et al. (2013)**, that the Southern Ocean provides ‘complete and uninterrupted connectivity’ between Antarctic and Southern Hemisphere killer whales. Despite the apparent homogeneity of the Southern Ocean it harbours geographically structured populations of many species and is a hotbed for adaptation (see examples given in **Rogers, 2007** and **Moon, Chown & Fraser, 2017**). For example, pelagic versus coastal niche, and oceanographic fronts, shape the range, dispersal potential and consequently genetic structuring among Southern Ocean penguin populations (**Clucas et al., 2018**). Our findings of structure between Antarctic and all other killer whales, in addition to previous findings of structure between Antarctic types *B1*, *B2* and *C* (**Foote et al., 2016**) are therefore consistent with patterns in other Antarctic taxa. Our findings make clear that, despite some connectivity, sub-Antarctic and Antarctic killer whale populations should not be conflated.

Ancient vicariance during a previous glacial cycle followed by more recent admixture is also inferred from the ancestry of the sympatric North Pacific mammal-eating *transient* ecotype and the fish-eating *resident* ecotype. The results from the PCA, PSMC and the *D*-statistics indicate more recent mean genome-wide coalescence and greater sharing of derived alleles and longer IBS tracts between the *resident* ecotype and most North Atlantic samples than between the *transient* ecotype and those same populations. We therefore infer that the *resident* ecotype shares a more recent common ancestor with these North Atlantic samples than does the *transient* ecotype. Our finding that alleles derived in the *transient* are shared more commonly with the *resident* than with all non-Pacific samples suggests gene-flow within the North Pacific between *residents* and *transients*, as first inferred from Isolation with Migration analyses (IMa) of microsatellite genotypes by **Hoelzel et al. (2007)**. Whilst this may appear to contradict long-term observations of social isolation between the two ecotypes (**Morton 1990**; **Baird & Dill, 1995**; **Ford, 2009**), gene flow via intermediary populations is supported by the *f*_4_-ratio test which identified Eastern Tropical Pacific samples and the *offshore* ecotype as having a mixture of *transient* and *resident* ancestry. Admixture may also be largely ancestral, rather than contemporary. The *f*_3_-statistic tests indicate greater drift relative to admixture in the *resident*, compared with the transient. In the PCA (Figure 1b), the segregation of samples along PC2 is driven by coalescence of shared genetic variation within the resident ecotype, *i.e*. lineage-specific drift in the *resident*. Our *resident* samples originate from across the ecotype’s North Pacific range, from Washington State, USA to the Sea of Okhotsk off Eastern Russia. Therefore, the variation segregating in the *resident* ecotype and driving PC2 in Figure 1b must pre-date the separation into the several *resident* sub-populations which have subsequently colonised much of the Pacific rim (**Filatova et al., 2018**). If this drift in allele frequencies shared among *residents* occurred post-admixture with the *transient* ecotype, it would increase genetic differentiation between the two currently sympatric North Pacific ecotypes. Identifying introgressed haplotype lengths will be an important next step in unravelling this detail of the evolutionary history of killer whale ecotypes.

This pattern of recurrent vicariance and subsequent admixture, likely corresponding to the cyclical expansion and contraction of high latitude habitat during the glacial cycles, contributes towards the genomic heterogeneity within an individual genome. Our results suggest that tracts originating from past vicariance during previous glacial cycles can be numerous and comprise a significant proportion of the genome. It is therefore important to consider such tracts in future analyses. For example, admixture between archaic hominin and modern Eurasian humans can inflate divergence time estimates between African and non-African populations (**Alves et al., 2012**). Similarly, the genomes of the Antarctic killer whales represent at least two different demographic histories: the major ancestry component reflects a history in which the Antarctic types appear to be recently derived from other Southern Ocean populations; the minor ancestry reflects an ancient divergence that predates the TMRCA of most other killer whale lineages. Thus, the previous estimated TMRCA of 126-227 KYA (**Foote et al., 2016**) will be an average of the variation in TRMCA across the genome, thereby ignoring the true complexity of vicariance and admixture among killer whale populations. Furthermore, depending upon the demographic and evolutionary history of these tracts, *e.g*. if they evolved in a locally adapted ancestral population or if the ancestral effective population size was small, they could harbour adaptive variation associated with this extreme Antarctic climate (as per **Racimo et al., 2015**) and/or weakly deleterious mutations (as per **Harris & Nielsen, 2016**; **Juric, Aeschbacher & Coop, 2016**). Thus, further research into the demographic history of these archaic tracts is warranted.

In summary, the global dataset of genomes analysed here contributes further to the emerging consensus (*e.g*. **Gopalakrishnan et al., 2018**; **Malinsky et al., 2018**; **Sinding et al., 2018; Tusso et al., 2018**) that the relationship among natural populations is rarely well represented as a bifurcating tree. The evolutionary history of natural populations can include episodic long-range dispersal, population replacement and admixture which greatly transform the distribution of global genetic variation. Furthermore, we highlight the importance of a phenomenon hitherto rarely considered in studies of non-human study organisms, that of archaic tracts within a genomic background with much younger TMRCA. Whilst past studies have simulated gene flow from unsampled ‘ghost’ populations (*e.g*. **Wilson & Bernatchez, 1998; Beerli, 2004**; **Slatkin, 2005**), by using tools previously primarily harnessed for the study of human population dynamics we highlight how genomic data can be leveraged to both identify the genomic regions derived from archaic and ghost populations and quantify their effect on contemporary population structure.

## Acknowledgements

The sequencing service was provided by the Genomics Core Facility (GCF), Norwegian University of Science and Technology (NTNU). GCF is funded by the Faculty of Medicine and Health Sciences at NTNU and Central Norway Regional Health Authority. We are grateful to the Museum of New Zealand, Te Papa Tongarewa, Wellington, for providing material from the type D specimen (#1077). A.D.F. was supported by a short visit grant from the European Science Foundation-Research Networking Programme ConGenOmics and by a Swisss National Science Foundation grant (31003A-143393) to L. Excoffier, and by the Welsh Government and Higher Education Funding Council for Wales through the Sêr Cymru National Research Network for Low Carbon, Energy and Environment, and from the European Union’s Horizon 2020 research and innovation programme under the Marie Skłodowska-Curie grant agreement No. 663830. We thank Alison Devault at Arbor Biosciences for support with the WGE experiment and Laurits Skov for advice on the HMM method.

