## Supplementary Materials for "Killer whale genomes reveal a complex history of recurrent admixture and vicariance"

### Table of Contents:


| | |
|---|---|
| Figure S1 | Page 2 |
| Figure S2 | Page 3 |
| Figure S3 | Page 4 |
| Figure S4 | Page 5 |
| Figure S5 | Page 5 |
| Figure S6 | Page 6 |
| Figure S7 | Page 7 |
| Figure S8 | Page 8 |
| Figure S9 | Page 9 |
| Figure S10 | Page 10 |
| Figure S11 | Page 11 |
| Figure S12 | Page 12 |
| Figure S13 | Page 13 |
| Figure S14 | Page 14 |
| Figure S15 | Page 15 |
| Figure S16 | Page 16 |
| Table S1 | Page 17 |
| References | Page 19 |

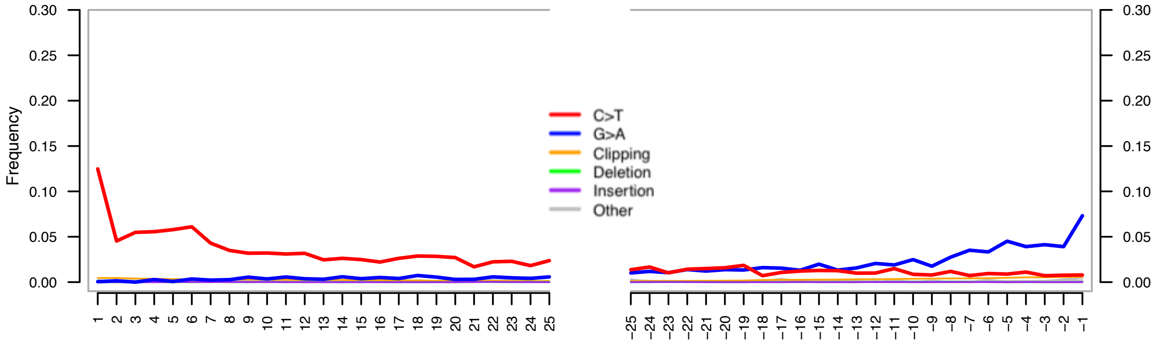


**Figure S1.** DNA misincorporation errors relative to the 5´ and 3´ read termini of a 41Mb scaffold (KB316842.1) of the genome of the *type D* killer whale museum specimen relative to the modern Norwegian killer whale reference. The two distributions for post mortem damage signatures (C>T and G>A) are shown in red and blue respectively. Nucleotide frequencies are shown for 25 bases upstream and downstream of the 5´ and 3´ read termini.

**(a)**

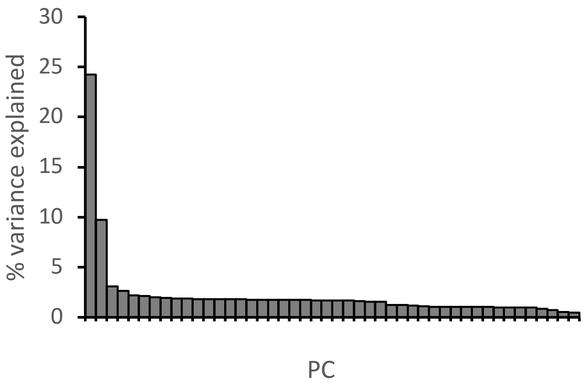


**(b)**

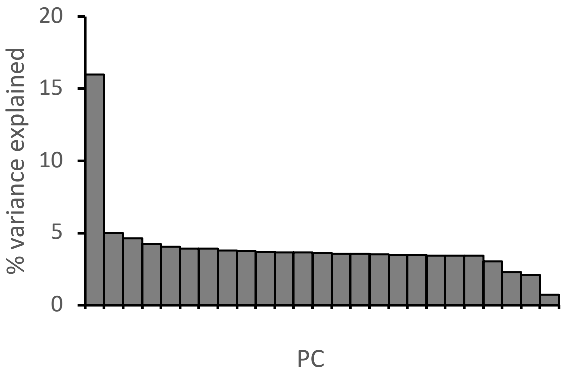


**Figure S2.** (a) Variance of the data explained by eigen vectors 1-46 of PCA in Figure 1b. (b) Variance of the data explained by eigen vectors 1-25 of PCA in Figure 1c.

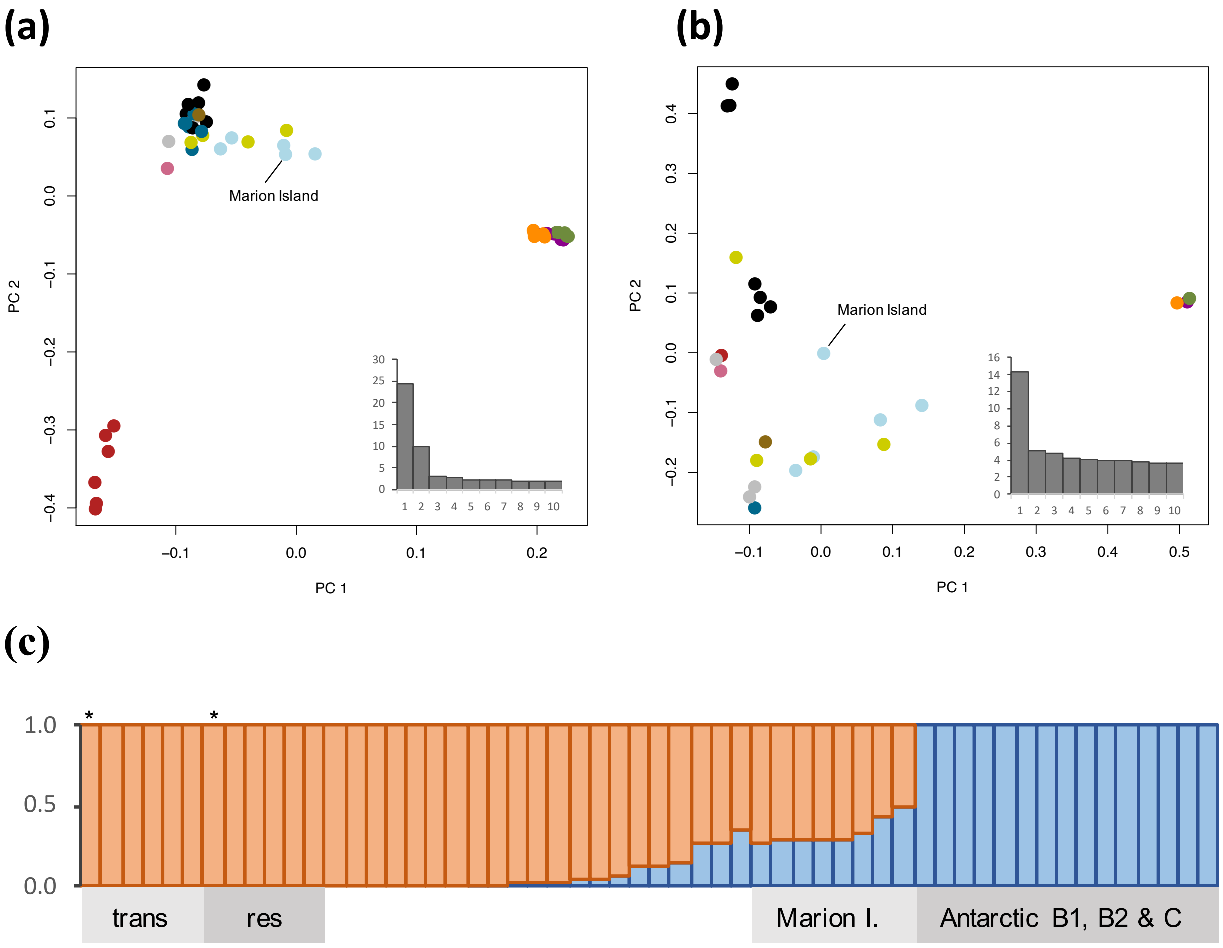


**Figure S3.** (a) PCA plots of the combined global and ecotype datasets, including RAD-seq data for a *resident*, *transient* and a subantarctic Marion Island killer whale. The RAD-seq samples for the *resident* and *transient* cluster with the whole genome sequence data indicating that batch effects between RAD and WGS data did not drive patterns observed in the PCA. Insert shows the variance of the data explained by eigen vectors 1-10. (b) PCA plot when only one 5× coverage genome per population and RAD-seq data for a subantarctic Marion Island killer whale are included. Insert shows the variance of the data explained by eigen vectors 1-10. In both (a) and (b) variation segregating in the Antarctic types *B1*, *B2* and *C* explains a significant ($P < 0.001$) component of the variation in the data. (c) Individual admixture proportions at $K = 2$, for WGS and RAD-seq data. *Transient* and *resident* RAD-seq samples are indicated by an asterisk, all eight Marion Island samples were from published RAD-seq data (Table S1). The analyses above were performed on 5,442 SNPs that were covered in all samples. In each analysis we see that some Southern hemisphere samples, including those from Marion Island, share some 'Antarctic' ancestry, but that there is clear structure separating the Antarctic types from these Southern populations.

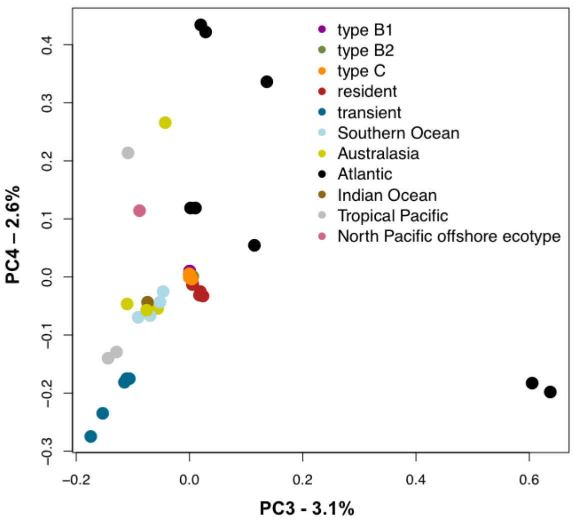


**Figure S4.** PCA plot of the combined global and ecotype datasets (46 individuals) showing PC3 and PC4. PC3 and PC4 are significant at $P < 0.001$ and $P < 0.05$ respectively.

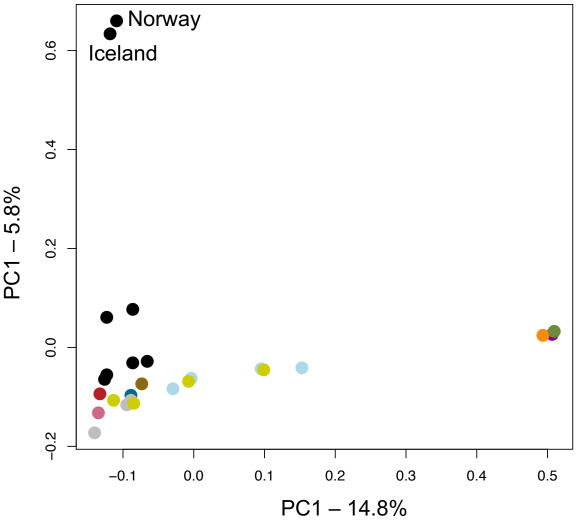


**Figure S5.** PCA plot in which one 5× coverage genome per population in included. The Iceland and Norway samples segregating along PC2, which is significant at $P < 0.001$.

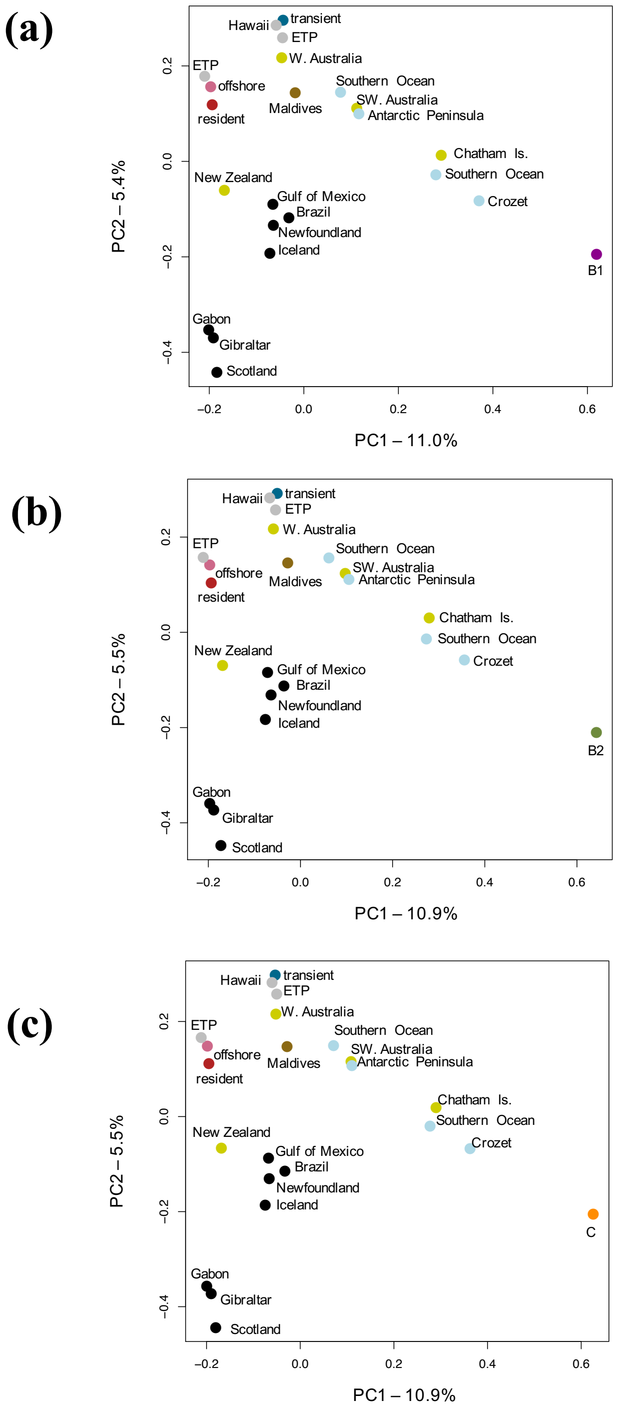


**Figure S6.** PCA plots in which one 5× coverage genome per population in included. Given their correlated allele frequencies, each PCA includes only one of the three Antarctic types: (a) *type B1*; (b), *type B2* and (c), *type C*. PC1 is significant ($P < 0.001$) in each case.

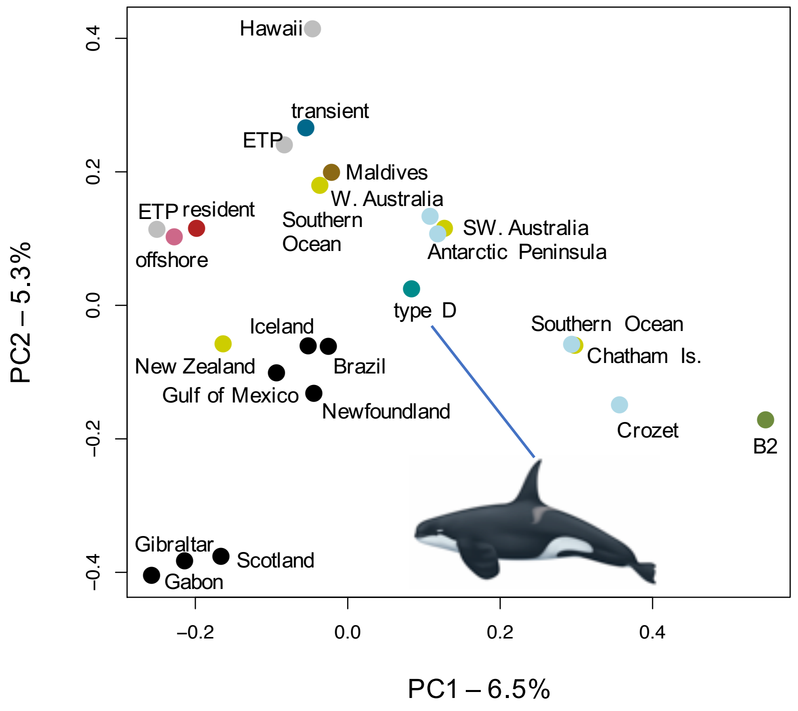


**Figure S7.** PCA plot including the *type D* killer whale morph, based on 6,565 transversions to avoid C>T misincorporations from DNA damage patterns in the genomic data generated from the *type D* museum sample.

**(a)**

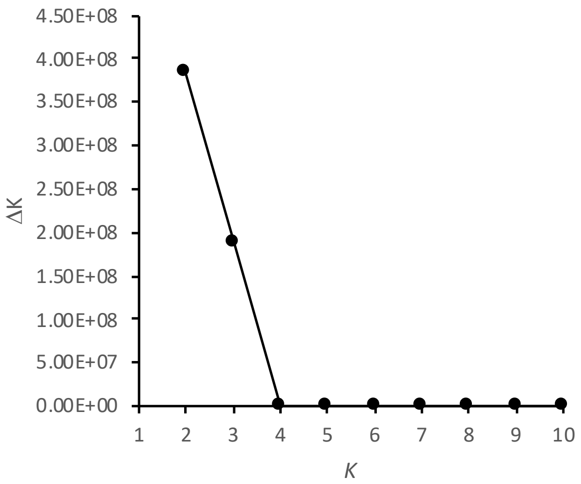


**(b)**

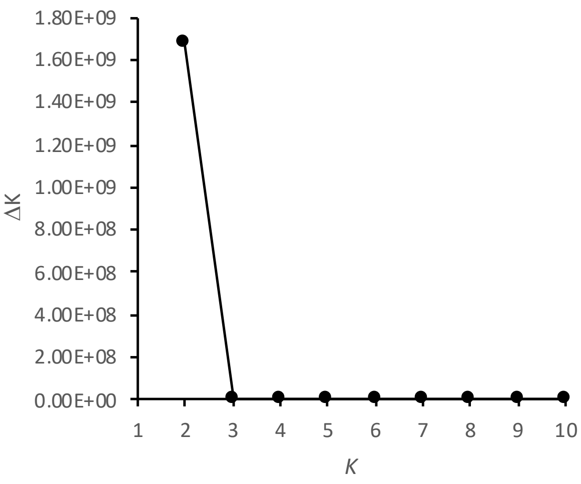


**Figure S8.** Mean values of $\Delta K$, the greatest step-wise increase in log likelihood (Evanno, Regnaut, & Goudet, 2005), from five runs at successive values of $K$ (number of genetic clusters). The uppermost hierarchical level of structure detected, based on $\Delta K$, is that for $K$=2 clusters. Antarctic types *B1*, *B2* and *C* form a single cluster and all other killer whales form another, with some 'Antarctic' ancestry detected in some southern hemisphere samples (see Figure 1e). For the dataset with multiple individuals from each ecotype (a), we observe a high increase in log likelihood between $K$=2 and $K$=3 clusters, in which the resident ecotype form a distinct cluster (see Figure 1d). (b) For the dataset with just one sample per population, the log likelihood increases at a steady rate above $K$=2.

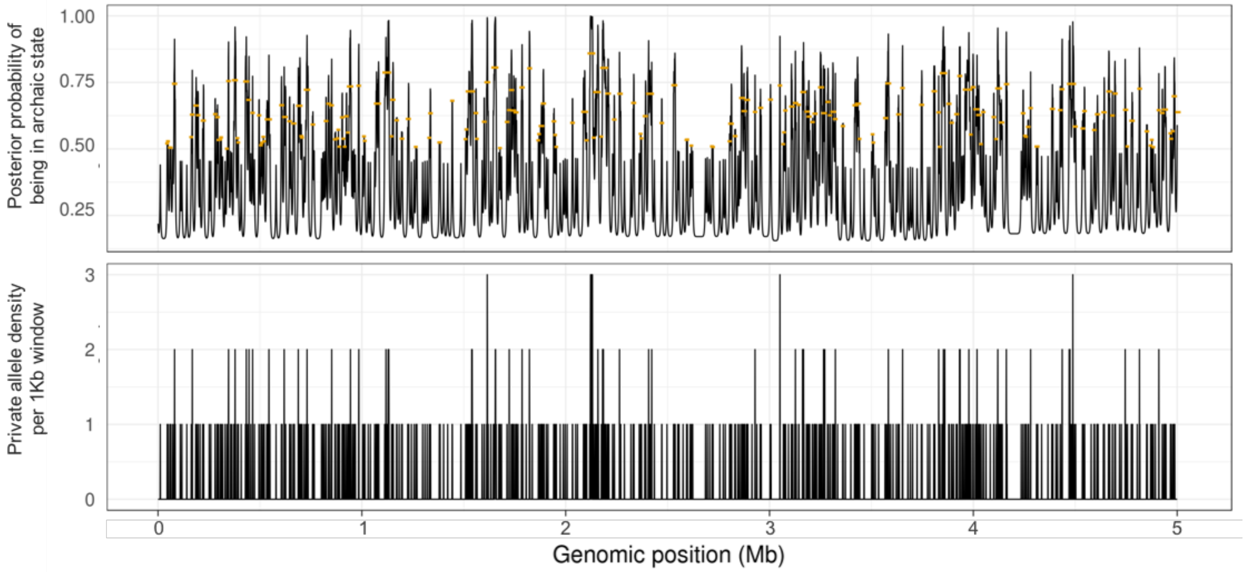


**Figure S9.** Top panel: The posterior probability of 1 kb windows being in the archaic state along 5 Mb of the genome of Antarctic *type B2*. The orange points are the mean posterior probability of tracts, which can be comprised of multiple windows, being in the archaic state. The signal is noisy and thus likely to be subject to false positives. The lower panel shows the density of SNPs per 1kb with alleles private to *type B2* when compared with the outgroup dataset.

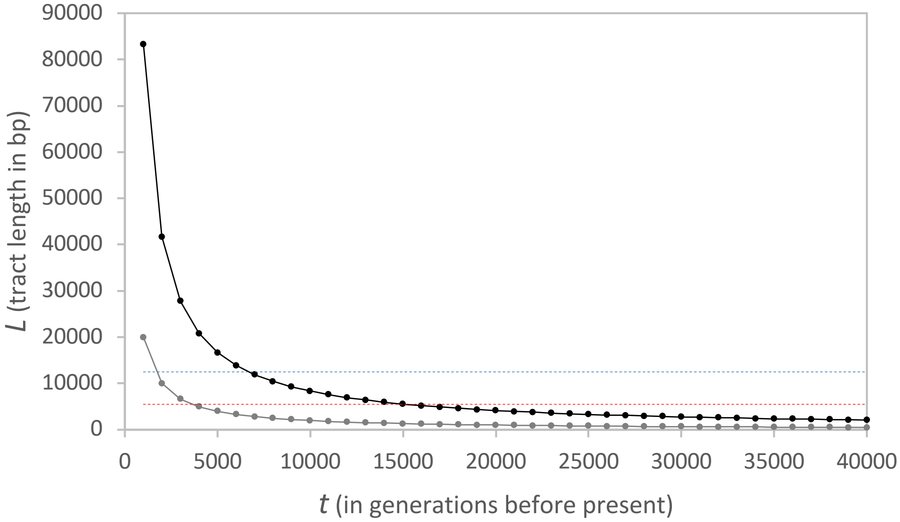


**Figure S10.** Expected age of ancestry tracts of different lengths, given a recombination rate ($r$) equivalent to that found in humans (as per Dumont & Payseur, 2008) of $1.2 \times 10^{-8}$/bp (1.2 cM/Mb) (black line), and a high recombination of $5.0 \times 10^{-8}$/bp (5.0 cM/Mb) (grey line). $L$ (expected tract length) is approximated as $1/(rt)$ (Racimo et al., 2015). Red dashed line marks the mean length (5,500 bp) of inferred archaic tracts with a posterior probability >0.5 in *type B2*. Blue dashed line marks the mean length (12,500 bp) of inferred archaic tracts with a posterior probability >0.8 in *type B2*.

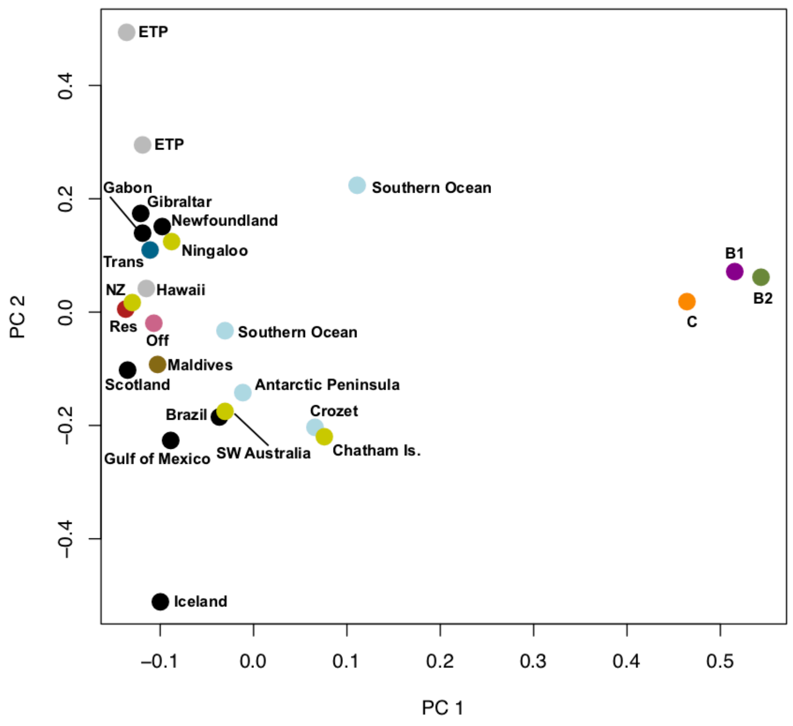


**Figure S11.** PCA plot based on 8,888 1 kb windows of the 41Mb autosomal scaffold (KB316842.1) inferred as archaic in the *type B2* killer whale. These regions contained a total of 25,208 SNPs, of which 4,406 were alleles private to *type B2* compared with the outgroup dataset used in the HMM. Note that due to linkage, these are not all independent observations. PC1 was the only significant component ($P > 0.001$) and explained of 18.3% of the variation in the data.

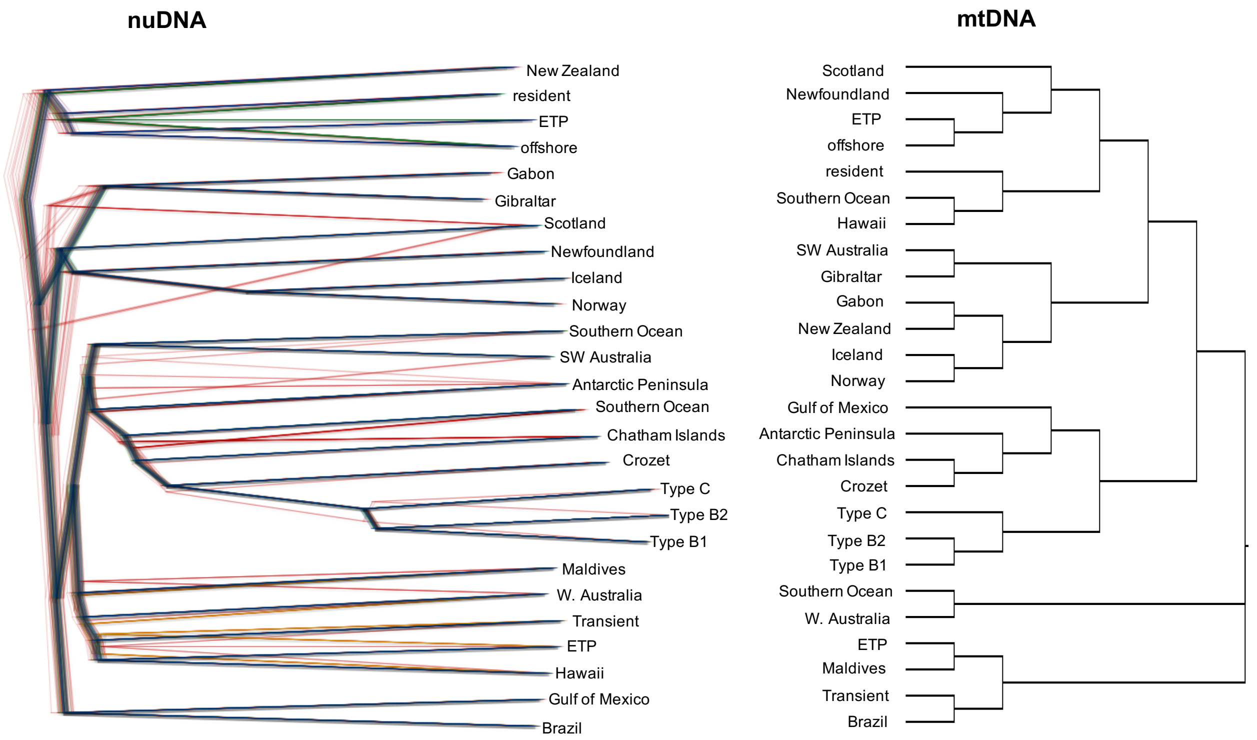


**Figure S12.** Maximum likelihood tree of mitogenomes (right) and tree reconstruction based on pairwise genetic distances between nuclear genomes (left). A tree cloud generated by DensiTree (Bouckaert, 2010) shows the diversity of trees representing 100 distance matrices generated by bootstrap sampling blocks of 500 SNPs from the original data set of 6,974,134 SNPs.

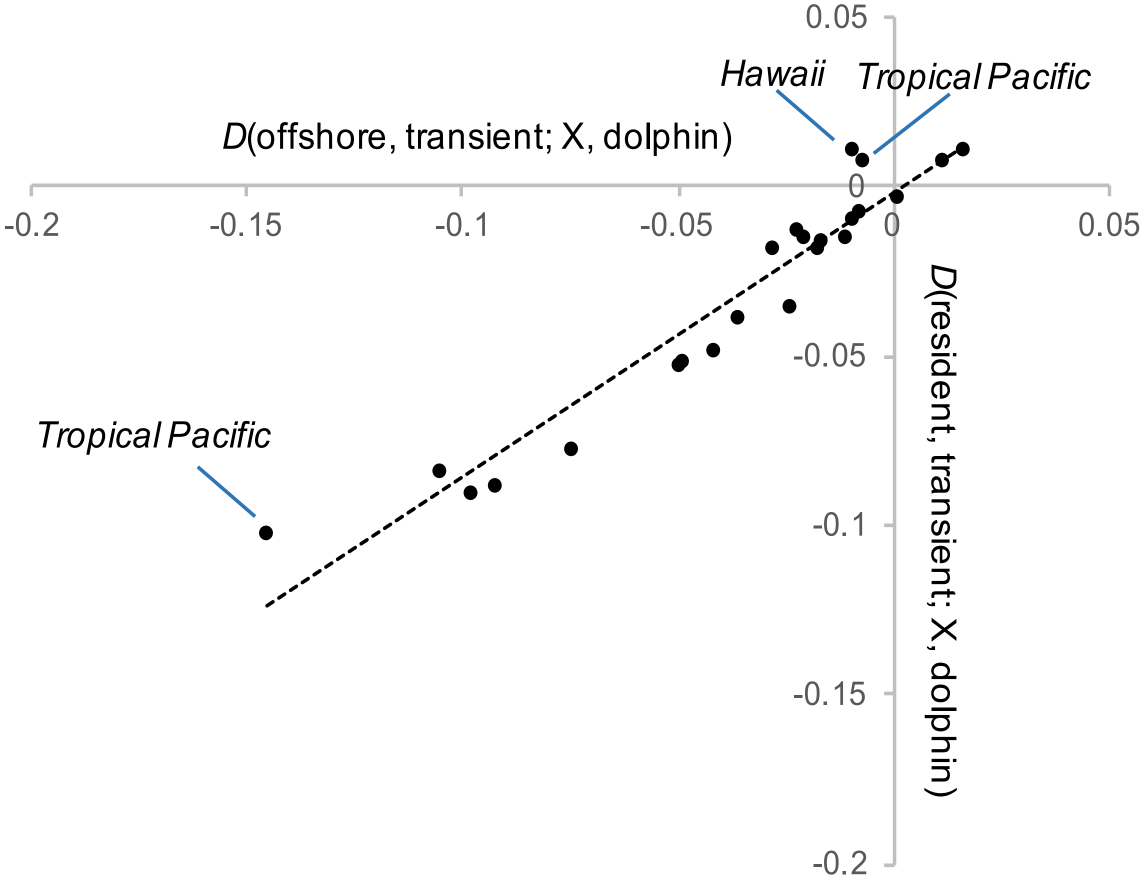


**Figure S13.** Estimates of D-statistics of the form *D*(offshore, transient; *X*, dolphin) and *D*(resident, transient; *X*, dolphin), where *X* represents a sample from the global dataset, are highly correlated (Pearson's correlation coefficient: $r_{23} = 0.9609$, $p < 0.00001$). Three moderate outliers (annotated) were either more negative along the *x*-axis due to a greater proportion of shared derived alleles between *X* and the offshore ecotype, or more positive along the *y*-axis due to a greater proportion of shared derived alleles between *X* and the *transient* ecotype, than predicted by the trend-line. The three individuals representing *X* in these outlier points were all sampled in the Tropical Pacific waters, and more recent gene flow between the Tropical Pacific populations and the *offshore* and *transient* ecotypes may be an explanatory factor as to why these values marginally stray from the trend line.

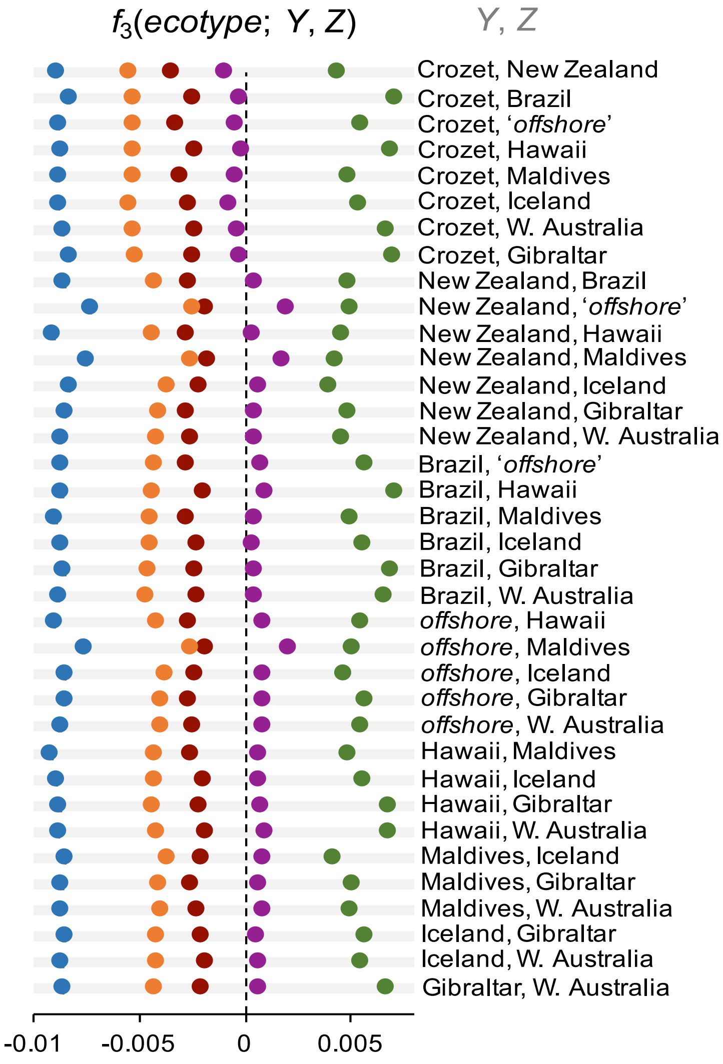


**Figure S14.** $F_3$-statistics of the form $f_3(X;\ Y, Z)$, where $X$ is one of the five focal ecotypes (blue = *transient*; red = *resident*; purple = *B1*; green = *B2*; orange = *C*), and $Y$ and $Z$ are potential source populations of ancestry in ecotype $X$. $F_3$-statistics were minimized and significantly ($Z$<-3) negative when estimating $f_3$(*transient*; $X$, $Y$) and were positively maximized when Antarctic *type B2* was the target ecotype for all tested combinations of $X$ and $Y$. $F_3$-statistics indicated the North Pacific *resident* ecotype and Antarctic *type C* are partly admixed with one or more of the donor populations, or some unsampled, but closely related populations (see for example, the outgroup case, **Patterson et al., 2012**). $F_3$-statistics were only significantly negative for *type B1* if the individual sampled off the subantarctic Crozet Archipelago was one of the donor populations ($X$ or $Y$).

**a,**

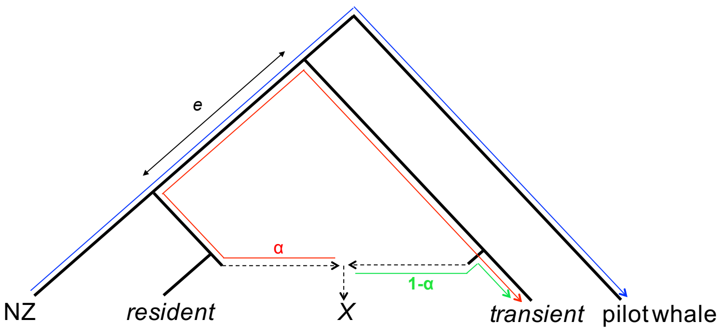


$f_4$(NZ,pilot whale;$X$,transient) / $f_4$(NZ,pilot whale;resident,transient) = α

**b,**

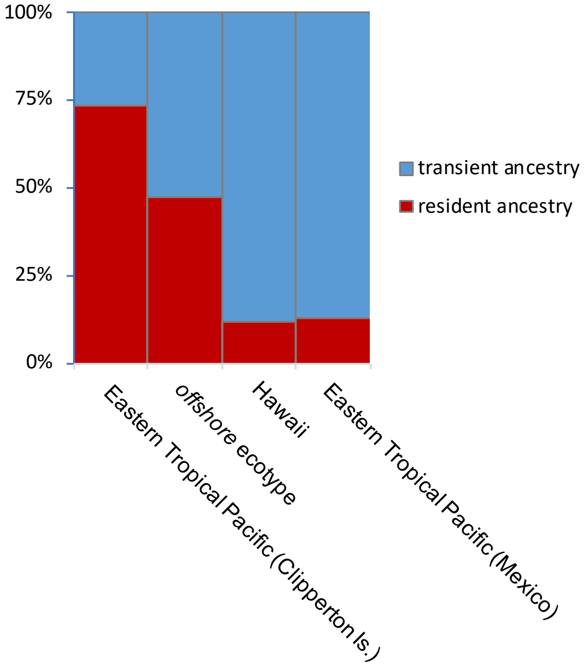


**Figure S15.** Relative ancestry proportions of North Pacific *resident* and *transient* ecotypes found in Tropical and North Pacific samples were estimating using **a,** $f_4$ ratio tests. **b,** The Eastern Tropical Pacific sample from Clipperton Island, close to Mexico, had the highest *resident* ancestry, consistent with the clustering patterns in other analyses in this study. The *offshore* sample shared approximately equal ancestry with the *resident* and *transient* ecotype. The Hawaiian and Mexican samples, shared mostly *transient* ancestry, consistent with their clustering with the *transient* ecotype in NGSadmix, ngsDist, D-statistic and PCA analyses.

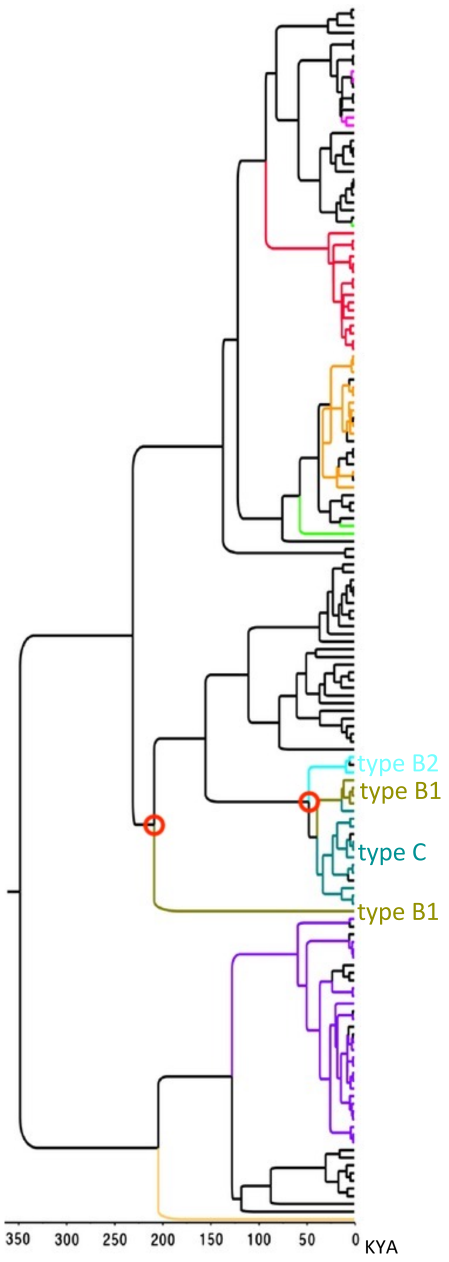


**Figure S16.** The TMRCA of types *B1*, *B2* and *C* based on mitochondrial genome sequences is estimated at approximately 50 KYA, with the exception of a unique mtDNA haplotype of a *type B1* individual from the Ross Sea, which had a TMRCA of over 200 KYA with the other haplotypes sampled from types *B1*, *B2* and C. This figure is adapted from **Morin et al., 2015**, see the original paper for full details of the mitogenome phylogeny generated from a comprehensive global dataset of 452 samples.

| Sample | Sequencing method | Population | Accession | Source |
|---|---|---|---|---|
| 28521 | WGS | offshore | TBC | This study |
| 106717 | WGS | Brazil | TBC | This study |
| 30525 | WGS | Hawaii | TBC | This study |
| 37927 | WGS | ETP | TBC | This study |
| 476 | WGS | Newfoundland | TBC | This study |
| 40918 | WGS | Maldives | TBC | This study |
| 92361 | WGS | Antarctic Peninsula | TBC | This study |
| 105084 | WGS | New Zealand | TBC | This study |
| 105085 | WGS | Chatham Isles | TBC | This study |
| 17221 | WGS | Southern Ocean | TBC | This study |
| 17222 | WGS | Southern Ocean | TBC | This study |
| 114343 | WGS | Gabon | TBC | This study |
| 11979 | WGS | ETP | TBC | This study |
| 39127 | WGS | Gulf of Mexico | TBC | This study |
| Gus | WGS | Ningaloo, W. Australia | TBC | This study |
| OO-160 | WGS | Strait of Gibraltar | TBC | This study |
| Urkel | WGS | Bremer Canyon, S.W. Australia | TBC | This study |
| Crozet | WGS | Crozet Archipelago | TBC | This study |
| GM6 (G6_04) | WGS | Pilot whale from Kerguelen | TBC | This study |
| M4/16 | WGS | Scotland | TBC | This study |
| 993 | WGS | Iceland | TBC | This study |
| AFOo | WGS | Norway | PRJNA167475 | Foote et al. 2015 |
| 94446 | WGS | transient | TBC | This study |
| 157515 | WGS | resident | SRP035610 | Moura et al. 2014 |
| 124038 | WGS | Antarctic type B1 | TBC | This study |
| 26619 | WGS | Antarctic type C | TBC | This study |
| 92365 | WGS | Antarctic type B2 | TBC | This study |
| 1077 | WGS | type D | TBC | This study |
| 73077 | WGS | Antarctic type B1 | ERS554425 | Foote et al. 2016 |
| 78580 | WGS | Antarctic type B1 | ERS554426 | Foote et al. 2016 |
| 88344 | WGS | Antarctic type B1 | ERS554430 | Foote et al. 2016 |
| 88348 | WGS | Antarctic type B1 | TBC | This study |
| 40882 | WGS | Antarctic type B2 | ERS554435 | Foote et al. 2016 |
| 32009 | WGS | Antarctic type B2 | ERS554434 | Foote et al. 2016 |
| 31884 | WGS | Antarctic type B2 | ERS554433 | Foote et al. 2016 |
| 92363 | WGS | Antarctic type B2 | ERS554439 | Foote et al. 2016 |
| 26614 | WGS | Antarctic type C | ERS554462 | Foote et al. 2016 |
| 26623 | WGS | Antarctic type C | ERS554466 | Foote et al. 2016 |

| | | | | |
|---|---|---|---|---|
| 26627 | WGS | Antarctic type C | ERS554468 | Foote et al. 2016 |
| 45804 | WGS | Antarctic type C | ERS554470 | Foote et al. 2016 |
| 126158 | WGS | resident | ERS554442 | Foote et al. 2016 |
| 126163 | WGS | resident | ERS554444 | Foote et al. 2016 |
| 126178 | WGS | resident | ERS554448 | Foote et al. 2016 |
| 35322 | WGS | resident | ERS554450 | Foote et al. 2016 |
| 57919 | WGS | transient | ERS554455 | Foote et al. 2016 |
| 79751 | WGS | transient | ERS554458 | Foote et al. 2016 |
| 79759 | WGS | transient | ERS554459 | Foote et al. 2016 |
| 62471 | WGS | transient | ERS554456 | Foote et al. 2016 |
| 032_AR | RAD-seq | resident | SRX701424 | Moura et al. 2015 |
| 047_AT | RAD-seq | transient | SRX701458 | Moura et al. 2015 |
| 131 | RAD-seq | Marion Island | SRX701576 | Moura et al. 2015 |
| 132 | RAD-seq | Marion Island | SRX701582 | Moura et al. 2015 |
| 135 | RAD-seq | Marion Island | SRX701586 | Moura et al. 2015 |
| 136 | RAD-seq | Marion Island | SRX701592 | Moura et al. 2015 |
| 137 | RAD-seq | Marion Island | SRX701598 | Moura et al. 2015 |
| 139 | RAD-seq | Marion Island | SRX701604 | Moura et al. 2015 |
| 142 | RAD-seq | Marion Island | SRX701610 | Moura et al. 2015 |
| 143 | RAD-seq | Marion Island | SRX701614 | Moura et al. 2015 |

**Table S1.** List of samples used in this study.
